# Lifelong dietary protein restriction accelerates skeletal muscle loss and reduces muscle fibre size by impairing proteostasis and mitochondrial homeostasis

**DOI:** 10.1101/2023.09.27.559536

**Authors:** Ufuk Ersoy, Moussira Alameddine, Ioannis Kanakis, Mandy Jayne Peffers, Malcolm J. Jackson, Katarzyna Goljanek-Whysall, Aphrodite Vasilaki, Susan E Ozanne, Gibran Pedraza Vazquez

## Abstract

The early life environment significantly affects the development of age-related skeletal muscle disorders. However, the long-term effects of lactational protein restriction on skeletal muscle are still poorly defined. Our study revealed that male mice nursed by dams fed a low-protein diet during lactation exhibited skeletal muscle growth restriction. This was associated with a dysregulation in the expression levels of genes related to the ribosome, mitochondria and skeletal muscle development. We reported that lifelong protein restriction accelerated loss of type-IIa muscle fibres and reduced muscle fibre size by impairing mitochondrial homeostasis and proteostasis at 18 months of age. However, feeding a normal-protein diet following lactational protein restriction prevented accelerated fibre loss and fibre size reduction in later life. These findings provide novel insight into the mechanisms by which lactational protein restriction hinders skeletal muscle growth and includes evidence that lifelong dietary protein restriction accelerated skeletal muscle loss in later life.

## Introduction

The nutritional environment experienced in early life can influence the risk of developing age-related diseases, including type 2 diabetes, cardiovascular diseases, and sarcopenia [1–4]. Maternal protein malnutrition is a concern in many populations globally, and it leads to permanent structural changes in organs, alterations in gene expression, and changes in cellular ageing in the offspring [1, 5–7]. It is widely accepted that maternal protein restriction during gestation limits the *in utero* growth of the offspring and has long-lasting effects that persist into later life through epigenetic modifications [1, 8, 9]. However, it is unclear whether maternal protein restriction during lactation protects the offspring through hormesis or has similar effects to maternal protein restriction during gestation [10, 11]. Reports from animal studies have demonstrated that lactational protein restriction impairs major metabolic pathways involved in the regulation of lifespan and metabolic syndrome in 21-day-old mice or in adult rats [12–14], but pups from dams on a low protein diet demonstrated an increased lifespan compared to control mice [15].

Muscle growth occurs through an increase in muscle fibre number or an increase in the size of individual fibres [16]. While postmitotic muscle fibres are formed during embryonic development via primary and secondary myogenesis [17], the early neonatal period in humans and rodents is also a critical time for skeletal muscle development as muscle hypertrophy mainly occurs during early postnatal life through a rapid increase in the number of myonuclei [18, 19]. Skeletal muscle is a highly dynamic tissue, and its development is particularly prone to nutritional deficiency compared with other tissues [20–22]. Suboptimal maternal nutrition during development dysregulated myogenesis, reduced the number of muscle fibres and muscle mass, and changed fibre type distribution in the offspring [22, 23]. Moreover, it has been shown that an isocaloric maternal low protein diet impacted skeletal muscle metabolism and mass, and the expression of key genes involved in mitochondrial metabolism in skeletal muscle of the offspring [12, 24–26]. Although lactational protein restriction reduced skeletal muscle weight in mice at weaning, it remains unclear whether these adverse effects persist in skeletal muscle following weaning onto a normal protein diet in later life [12].

In this study, we investigated the effects of lactational protein restriction on global gene expression changes in skeletal muscle at weaning and identified that genes involved in ribosomal homeostasis, mitochondrial function, myofibre and muscle development were affected. We further demonstrated that lifelong feeding a low-protein diet after lactational protein restriction accelerated type-IIa fibre loss and reduced muscle fibre size at 18 months of age by dysregulating ribosomal gene expression, autophagy, AMP-activated protein kinase (AMPK) signalling, and mitochondrial homeostasis. Some of the adverse effects of maternal protein restriction during lactation on skeletal muscle were corrected following weaning onto a normal protein diet in 18-month-old male mice.

## Results

### Maternal protein restriction during lactation reversibly inhibited growth in their offspring

To study the effects of lactational protein restriction, control mice were nursed either by dams fed a low-protein diet (8%) (NL) or a normal protein content diet (20%) (NN) during lactation (Fig. 1a). As expected, the NL mice exhibited significantly smaller body weight, as well as lower gastrocnemius (GAS) muscle weight in comparison to the control group (NN) at day 21 (Fig. 1b, c). Lower body weight in early life is a predictor of skeletal muscle disorders in later life [2]. Therefore, we investigated whether the reduction in body weight and muscle weight persists or could be corrected by weaning onto a normal protein diet. By 3 months of age, the group of NL mice that were maintained on a normal protein diet (NLN) from weaning exhibited catch-up growth as indicated by body weight, GAS and soleus (SOL) muscle weights (Fig. 1b, d, e). Moreover, NLN mice had similar body weight and muscle weights at 18 months of age compared to the control (NNN) mice (Fig. 1b, f, g). In order to investigate the effect of lifelong protein restriction on skeletal muscle, we fed NL mice with a low protein diet (NLL) following weaning until the end of the study at 3 or 18 months of age. NLL mice exhibited significantly lower body weight and reduced SOL muscle weight compared to the control group (NNN), at both 3 and 18 months of age (Fig. 1b, e, g). Additionally, GAS muscle weight was significantly lower in 3-month-old NLL mice compared to the control group (Fig. 1d), but not at 18 months of age (Fig. 1g).

**Fig. 1.**
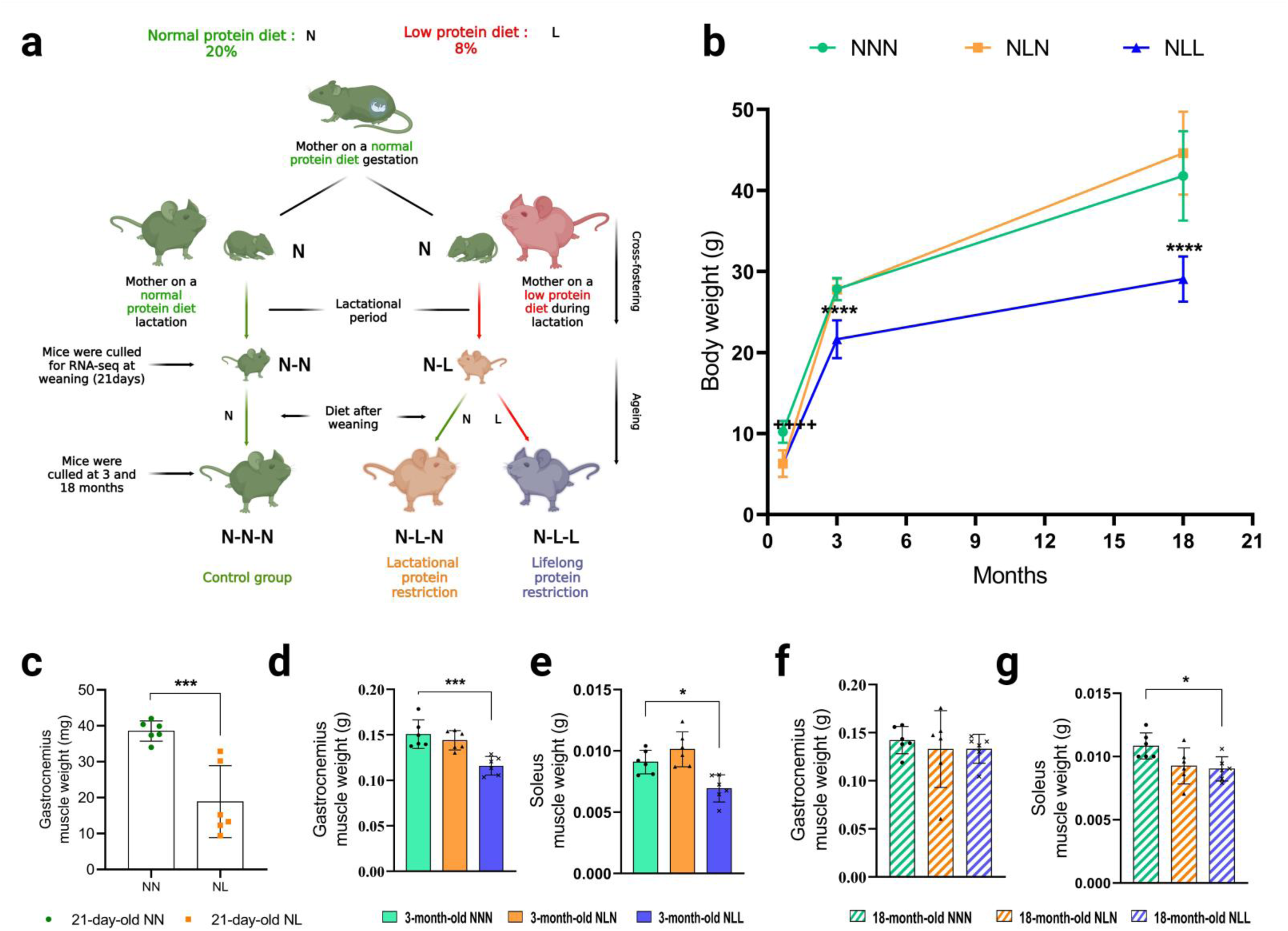
Illustrative description of experimental design, body weight and muscle weight. (**a**) Normal-Normal-Normal (NNN) litters received a normal protein diet during gestation and postnatally. Normal-Low(NL)-Normal (NLN) litters nursed by low-protein-fed dams during lactation and received a normal protein diet after weaning. Normal-Low(NL)-Low (NLL) litters were nursed by low-protein-fed dams during lactation and kept on a low-protein diet after weaning. **(b)** Body weights (g) of NNN, NLN, and NNN mice at 21-day-old, 3-month-old, and 18-month-old. *n=*6. **(c)** GAS muscle weights (g) of NN and NL mice at 21-day-old. *n=*6. **(d, e)** GAS muscle weights (g) of NNN, NLN, and NLL mice at (d) 3-month-old, (e) 18-month-old of age. *n=*6. **(f, g)** SOL muscle weights (g) of NNN, NLN, and NLL mice at (d) 3-month-old, (f) 18-month-old of age. *n=*6. Results are expressed as the mean ± standard deviation (mean ± SD). ⃰ ++++ p<0.0001 shows the significant difference between NN and NL mice in figure 1b. ⃰ p<0.05, ⃰ ⃰ p<0.01, ⃰ ⃰ ⃰ p<0.001, ⃰⃰⃰ ⃰ ⃰ ⃰ p<0.0001 shows the significant difference between NNN and NLL mice. Statistical comparisons were performed using ordinary one-way ANOVA with a Dunnett’s multiple comparisons test, considering NNN as the control group.

### RNA sequencing identified differences in the expression of key pathways associated with growth restriction in 21-day-old lactational protein-restricted mice

To reveal the underlying mechanisms of protein restriction during lactation, we compared the transcriptomic profiles of GAS muscles from 21-day-old NL and NN mice using RNA sequencing (RNA-seq) analysis. Of the 36048 mouse genes, between 43.4% and 54.2% had at least one read aligned; 13386 of the genes had no reads aligned from any of the 12 samples. 18246 protein coding genes and 4304 non-protein coding genes were found. Principal-component analysis (PCA) demonstrated that replicates from NN mice clustered together, while NL mice exhibited a more dispersed pattern along two principal components PC1 (37% of the total variance) and PC2 (13% of the total variance) (Fig. 2c). Analysis of differentially expressed genes demonstrated that expression of 257 genes was significantly increased (fold change>1.3 and adjusted p-values<0.1) and 282 genes were significantly reduced (fold change<-1.3 and adjusted p-values<0.1) in GAS muscle from NL mice compared to NN mice (Fig. 2d and Supplementary Data 1). Less stringent filters were chosen in order to capture a broader range of expression levels and these results were further validated. Differentially expressed genes were also filtered based on a |fold change|<2 and adjusted p-values<0.1, and the list of these genes can be found in Supplementary Data 1.

**Fig. 2.**
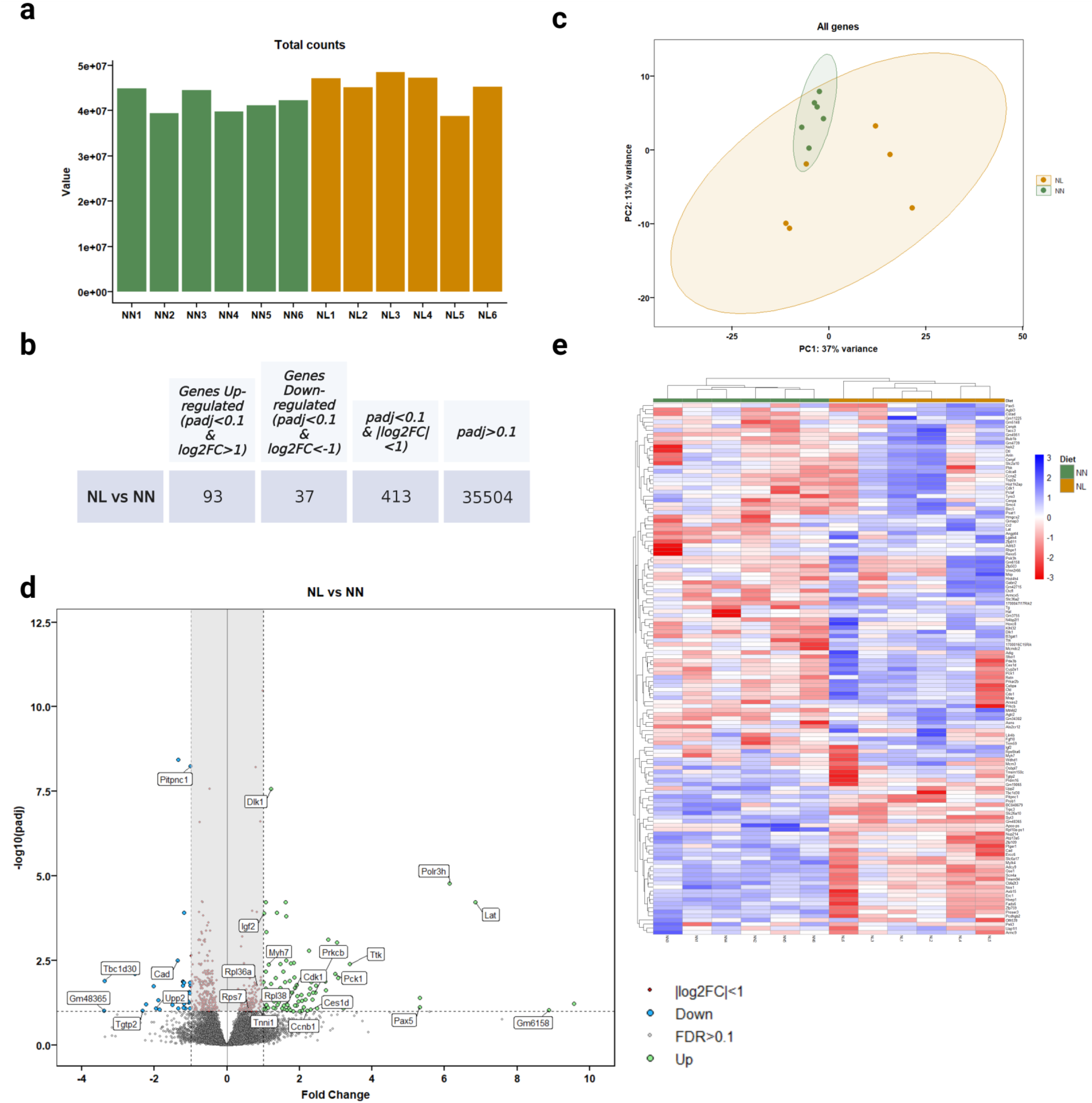
Identification of genes associated with the growth restriction phenotype seen in 21-day-old lactational protein-restricted mice. (**a**) Total count number of NN and NL samples. *n*=6. **(b)** Gene count of differentially expressed genes in skeletal muscle from NL vs NN mice**. (c)** Principal component analysis indicated NN mice were grouped together, whilst NL mice exhibited a spread pattern along two principal components PC1 (37% of the total variance) and PC2 (13% of the total variance). *n*=6. **(d)** Volcano plot summarising upregulated (green) and downregulated (blue) genes |Log2FC|>1 and adjusted p-values<0.1 in NL mice. Genes with adjusted p-values<0.1 and |Log2FC|<1 are highlighted on the middle of the plot and illustrated with red dots. **(e)** Heatmap summarising expression levels of upregulated or downregulated genes (|Log2FC|>1 and adjusted p-values<0.1) in groups. The gene expression level is presented as the read count. The colour code scale indicates the normalized counts (ranging from −3 (red) up to 3 (blue)).

Functional annotation of the upregulated genes revealed that these genes significantly associated with biological processes related to ribosomes and cell cycle (Fig. 3a and Supplementary Table 2). 29 large ribosomal subunit-encoding genes and 22 small ribosomal subunit-encoding genes were significantly upregulated in GAS muscle from NL mice (Supplementary Fig. 2). Notably, our findings revealed that the regulation of mitochondrial ATP synthesis coupled electron transport (enrichment false discovery rate (FDR)= 0.0008), the transition between fast and slow fibre (enrichment FDR=0.0002), and ribosomal small subunit assembly (enrichment FDR<0.0001) were amongst the highest upregulated Gene Ontology (GO) biological processes based on enrichment FDR (Fig. 3b and Supplementary Table 3). These data suggest that lactational protein restriction dysregulated the expression of genes involved in ribosomal homeostasis, mitochondrial homeostasis and skeletal muscle fibre development and distribution.

**Fig. 3.**
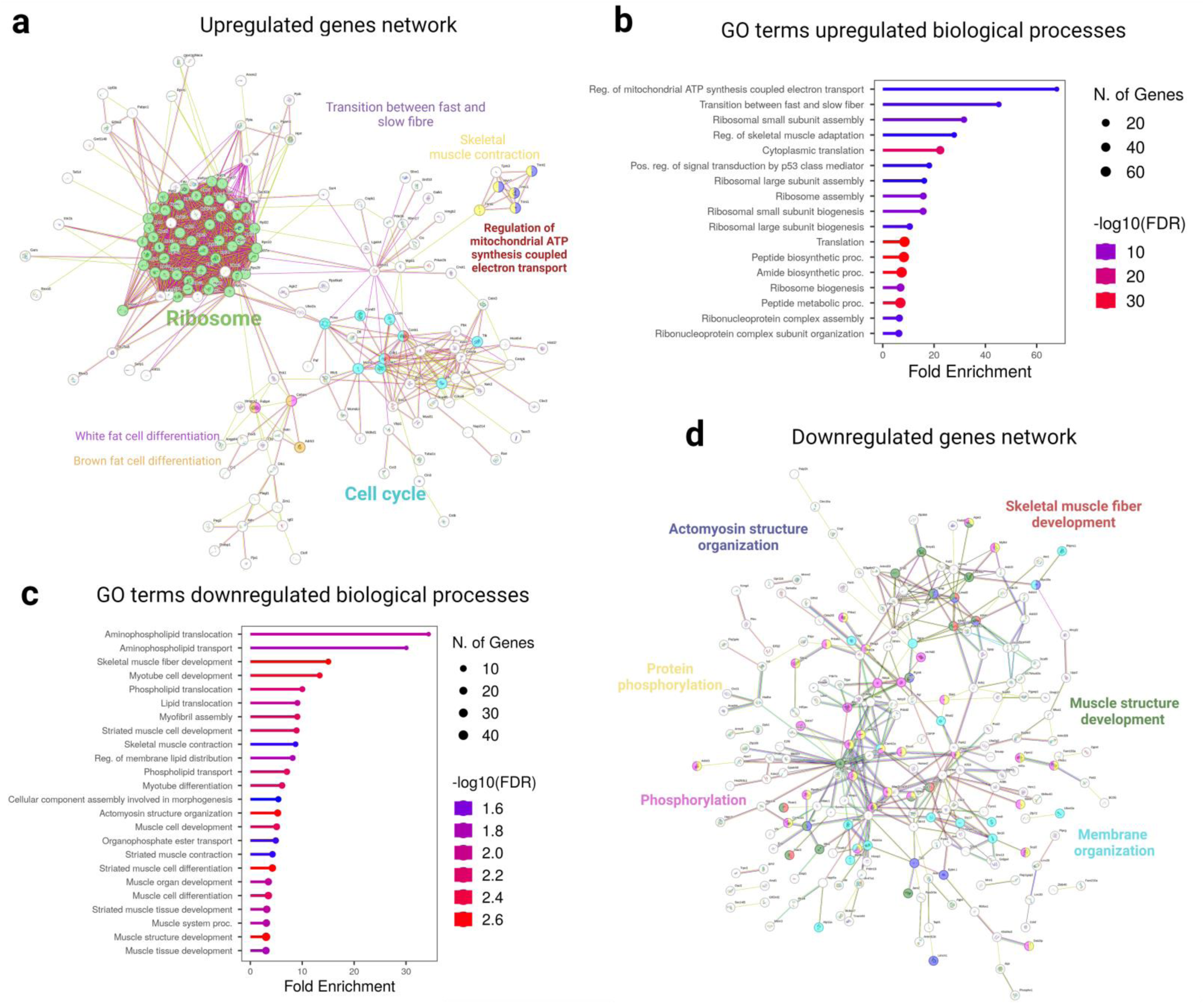
Identification of pathways associated with the growth restriction phenotype seen in 21-day-old lactational protein-restricted mice. (**a**) Bioinformatic network of significantly upregulated genes in GAS skeletal muscle from lactational protein restricted mice offspring using STRING. **(b)** The Gene ontology (GO) terms associated with upregulated genes. Analysis of the GO terms indicate that the most significantly fold-enriched biological processes include the regulation of mitochondrial ATP synthesis coupled with electron transport, the transition between fast and slow fibre types, the assembly of ribosomal small subunits, and the regulation of skeletal muscle adaptation. **(c)** The Gene ontology (GO) terms associated with downregulated genes. Based on the analysis of GO terms, the most significantly fold-enriched biological processes include the skeletal muscle fibre development, myotube cell development, myofibril assembly, skeletal muscle contraction, actomyosin structure organisation, and phosphorylation. **(d)** Bioinformatic network of significantly downregulated genes in GAS skeletal muscle from lactational protein restricted mice offspring using STRING.

To identify the pathways downregulated by lactational protein restriction, we analysed the supressed genes. Strikingly, multiple genes were associated with skeletal muscle fibre development (enrichment FDR=0.002), actomyosin structure organisation (FDR=0.002), phosphorylation (FDR=0.002), protein phosphorylation (FDR=0.006), myotube cell development (FDR=0.003), skeletal muscle contraction (FDR=0.02), myotube differentiation (enrichment FDR=0.006), and muscle cell development (FDR=0.005) (Fig. 3c, 3d and Supplementary table 5). Notably, mitophagy (FDR=0.046) and calcium signalling (FDR=0.046) were enriched (Supplementary table 4).

Therefore, we further assessed ribosomal gene expression, autophagy markers, AMPK signalling, mitochondrial homeostasis, skeletal muscle fibre number, muscle fibre size and fibre type distribution in NLL and NLN mice at 3 months or 18 months of age. Samples from 3-month and 18-month old mice were not sequenced.

### Lifelong protein restriction reduced total fibre number and fibre size, and remodelled muscle fibre type distribution

Since skeletal muscle development was disturbed in 21-day-old NL mice and genes involved in the transition between fast and slow fibre type were downregulated, we hypothesised that feeding a lifelong low-protein diet following lactational protein restriction would accelerate skeletal muscle fibre loss and remodel fibre type distribution. To confirm this, we first investigated the gene expression levels of myosin heavy chain (Myh) isoforms and demonstrated that only the expression level of Myh type-IIa was significantly reduced in SOL muscle from 3-month-old NLL mice compared to the control mice (Fig. 4a). We found significantly less fibres in the SOL muscle from 18-month-old NLL mice than in the control group (Fig. 4b, 4f). Additionally, minimum Feret’s diameter analysis showed a trend for a decrease in large muscle fibres (50.1-60 and 60+threshold), while the percentage of medium size fibre (30.1-40 threshold) was significantly increased in SOL muscle (Fig. 4d). Interestingly, only the number of type-IIa fibres significantly decreased in SOL, while the number of type-I and type-IIx/b fibres were similar to the control mice at 18 months of age (Fig. 4b). Moreover, type-IIa fibre type proportion (Fig. 4g) was significantly reduced in SOL from 18-month-old NLL mice, but fibre type proportion of type-I (Fig. 4h) and type-IIx/b (Fig. 4i) fibres were similar in the three different groups at 18 months of age.

**Fig. 4.**
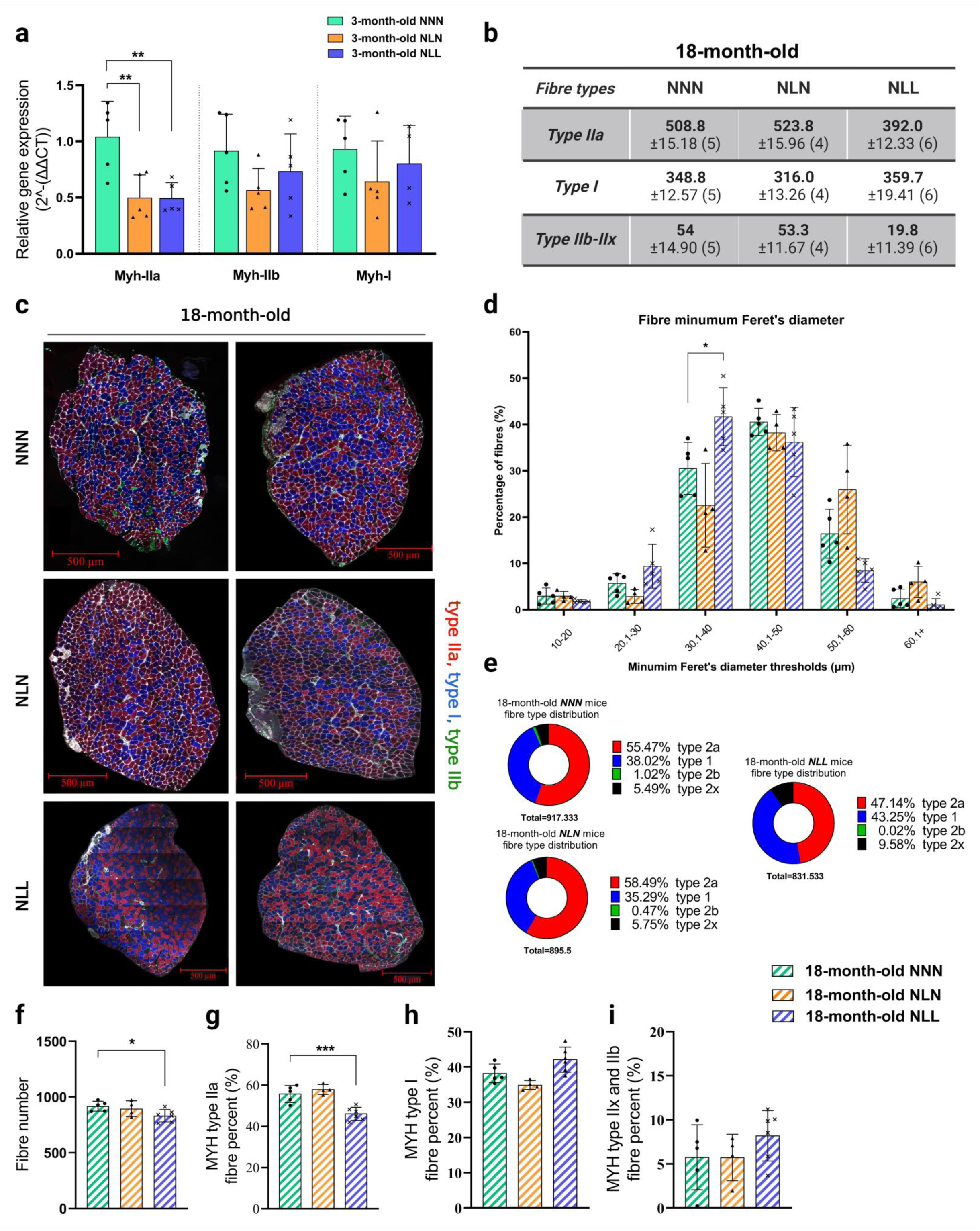
Lifelong protein restriction reduces total fibre number and muscle fibre size, and remodels muscle fibre type distribution. (**a**) qPCR analysis of Myosin Heavy Chain (Myh) isoforms in SOL muscle from 3-month-old NNN, NLN, and NLL mice. **(b)** The table shows the total number of different fibre types in 18-month-old NNN, NLN, and NLL mice. The total number of type-IIa fibres was reduced in NLL compared to the control group at 18-month-old of age. *n=4-*6. **(c)** Representative confocal images of MYH type-IIa (red), MYH type-I (blue, adjusted from orange), MYH type-IIb (green), and MYH type-IIx (unstained, black) in SOL muscle from 18-month-old mice. Scale bar, 500 µm. *n=4-*6. **(d)** Muscle fibre minimum Feret’s diameter. *n=4-*6 **(e, h, g, i)** The illustrations demonstrate the percentages of different MYH type-IIa (h), MYH type I (g), and MYH type-IIb and IIx (i) in SOL muscle from 18-month-old mice. *n=4-*6. **(f)** Total fibre numbers of SOL muscle from different mice. *n=4-*6. ⃰ p<0.05, ⃰ ⃰ p<0.01, ⃰ ⃰ ⃰ p<0.001, ordinary one-way ANOVA with a Dunnett’s multiple comparisons test.

We next assessed the effects of weaning onto a normal protein diet after lactational protein restriction on skeletal muscle in SOL from 18-month-old NLN mice. There was no difference in skeletal muscle fibre number between NLN and the control mice (Fig. 4f). Additionally, fibre size (Fig. 4d), fibre type distribution (Fig. 4e) and percentage (Fig. 4g, 4h, 4i) in NLN mice were similar to the control group.

Since the loss of muscle fibres primarily occurred from fast twitch fibres in the SOL muscle, we investigated the mechanistic changes in the GAS muscle, which consists of predominantly type II fibres.

### Lifelong protein restriction disturbed ribosomal gene expression and autophagy in adulthood and later life

We next validated the three most upregulated ribosomal genes; ribosomal protein L36A (Rpl36a), ribosomal protein L38 (Rpl38) and ribosomal protein S7 (Rps7) in GAS muscle from 21-day-old, 3-month-old, and 18-month-old mice. Similar to the RNA-seq data, significant increases in the expression levels of Rpl36a, Rpl38, and Rps7 were observed in NL mice at 21 days of age (Fig. 5a). This might be a compensatory mechanism to the lactational protein restriction-induced growth restriction. Unlike 21-day-old mice, there was no difference in Rpl36a, Rpl38, and Rps7 relative gene expression between NNN and NLL mice at 3 months of age (Fig. 5b). However, Rpl36a and Rpl48 were significantly downregulated in 18-month-old NLL mice compared to the control group (Fig. 5c). p62 (Sequestosome-1) is an autophagosome protein required for the formation and degradation of protein bodies containing Microtubule-associated proteins 1A/1B light chain (LC3), which accumulates when the autophagy is inhibited [27]. The amount of LC3-II present is closely associated with the number of autophagosomes formed; therefore, LC3 immunoblotting serves as a good indicator of autophagosome formation [28]. We found a significant reduction in p62 protein abundance and slight increase in the LC3B-II/LC3B-I ratio in skeletal muscle from NLL mice at 3 months of age, suggesting induced autophagy (Fig. 5d, 5e). This may account for the decrease in growth (Fig. 1b) and the reduced muscle weight (Fig. 1d, 1e) observed in NLL mice in adulthood.

**Fig. 5.**
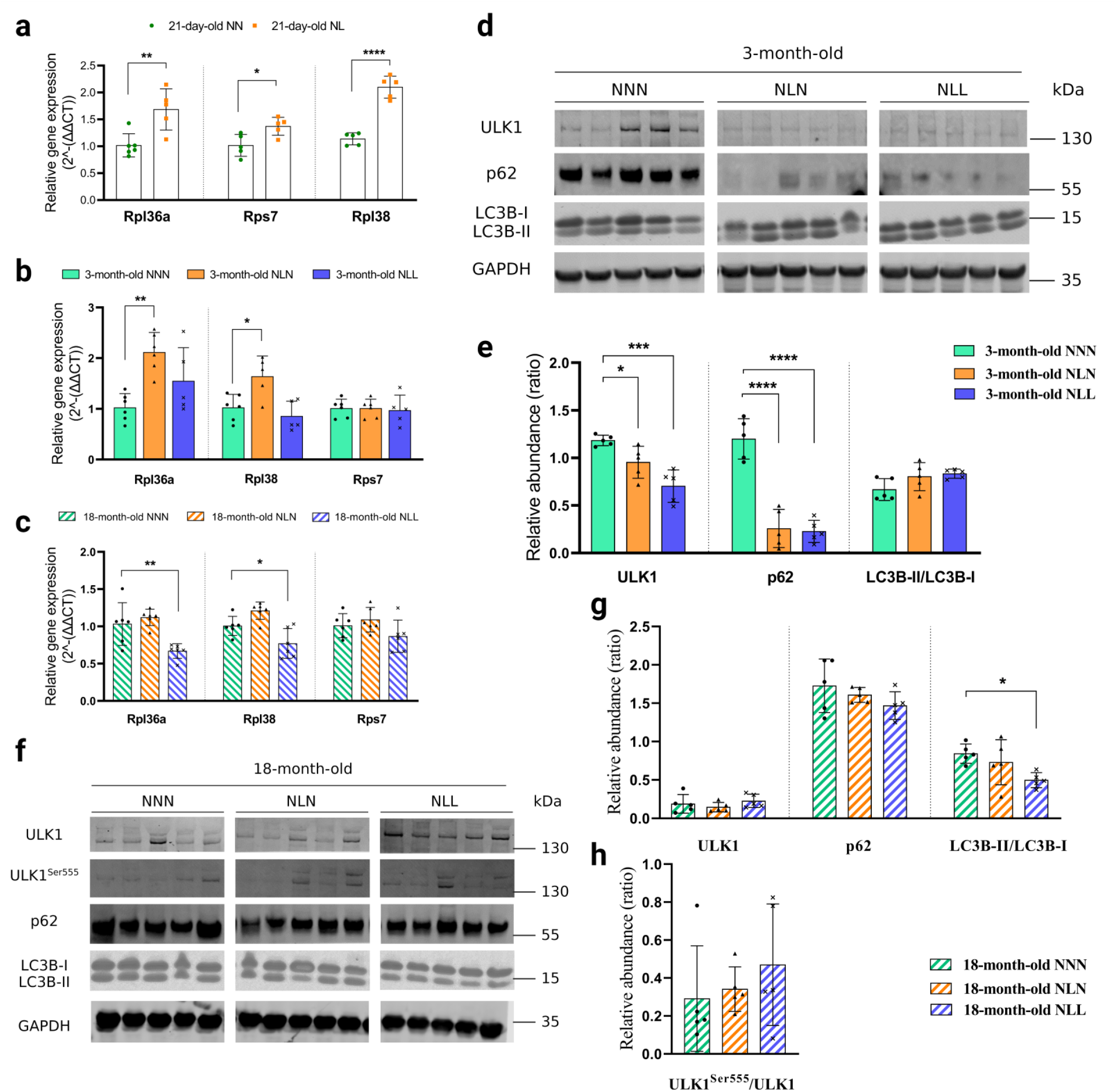
Lifelong protein restriction disturbs ribosomal gene expressions and autophagy in later life. (**a, b, c**) Relative gene expression levels of in (a) ribosomal protein L36A (Rpl36a), (b) ribosomal protein L38 (Rpl38), and (c) ribosomal protein S7 (Rps7) GAS muscle from 3-month-old mice *n=5-*6. **(d, e)** (d) Western blot analysis of UNC-51-like kinase 1 (ULK1), p62 (Sequestosome-1), and Microtubule-associated proteins 1A/1B light chain 3B (LC3B) in GAS muscle from 3-month-old NNN, NLN, and NLL mice. Samples were run on the same gel and images were cropped only for the purpose of this figure. GAPDH or ponceau staining were used as loading control. Source data provided as a Source data file. *n=5.* (e) Quantification of ULK1, p62, and LC3B-II/LC3B-I protein abundance levels. **(f, g)** (f) Western blot analysis of ULK1, p62, and LC3B in GAS muscle from 18-month-old NNN, NLN, and NLL mice Samples were run on the same gel and images were cropped only for the purpose of this figure. GAPDH was run on different gel and used as loading control. Ponceau staining was also applied to check equal loading. Source data provided as a Source data file. *n=5.* (g) Quantification of ULK1, p62, and LC3B-II/LC3B-I protein abundance levels*. n=5.* **(h)** Ratio of ULK1^Ser55^/ULK1 protein abundance. *n=5.* ⃰ p<0.05, ⃰ ⃰ p<0.01, ⃰ ⃰ ⃰ p<0.001, ⃰⃰⃰ ⃰ ⃰ ⃰ p<0.0001, ordinary one-way ANOVA with a Dunnett’s multiple comparisons test

While the LC3B-II/LC3B-I ratio was significantly reduced in NLL mice compared with the control group, p62 protein abundance was similar between groups at 18 months of age. Contrary to the observations in 3-month-old mice, these results indicate that feeding a low-protein diet following lactational protein restriction supressed autophagy in skeletal muscle in later life.

### Lifelong protein restriction impaired AMPK signalling and mitochondrial homeostasis in skeletal muscle

AMPK acts as a central regulator of mitochondrial homeostasis by modifying numerous aspects of the mitochondrial life cycle, including biogenesis, dynamics, and mitophagy [29, 30]. We hypothesised that the changes in expression levels of mitochondria-related genes and significant reduction in Prkab2 (Protein Kinase AMP-Activated Non-Catalytic Subunit Beta 2) expression level (Supplementary Data 1) in 21-day-old NL mice had further effects on AMPK signalling and mitochondrial homeostasis in NLL mice in adulthood or later life. To this end, we investigated the AMPK signalling pathway in NLL mice which were maintained on a low-protein diet following weaning. There was a significant reduction in AMPKα relative gene expression level (Fig. 6a) and protein content (Fig. 6b, 6c) in 3-month-old NLL mice, compared to control group. Interestingly, we found an increase in the ratio of AMPKα^Thr172^/AMPKα protein abundance in these mice (Fig. 6d), indicating an enhanced energy stress level in their GAS muscle. Our findings from 18-month-old NLL mice showed the reduction in AMPKα at the protein level was maintained, while there was no difference in AMPKα gene expression level (Fig. 7a) and AMPKα^Thr172^/AMPKα ratio (Fig. 7f). These results support that there was an increased energy stress level in GAS muscle from NLL mice. We assessed the mitochondrial content by investigating the activity and abundance of citrate synthase in GAS muscles. There was a significant reduction in both activity and abundance of citrate synthase in NLL mice at 3 months (Fig. 6e, 6f, 6g, 6h) and 18 months of age (Fig. 7b, 7c, 7e). Together, these findings suggest that mitochondrial biogenesis and mitochondrial density were reduced in the GAS muscle from NLL mice [31].

**Fig. 6.**
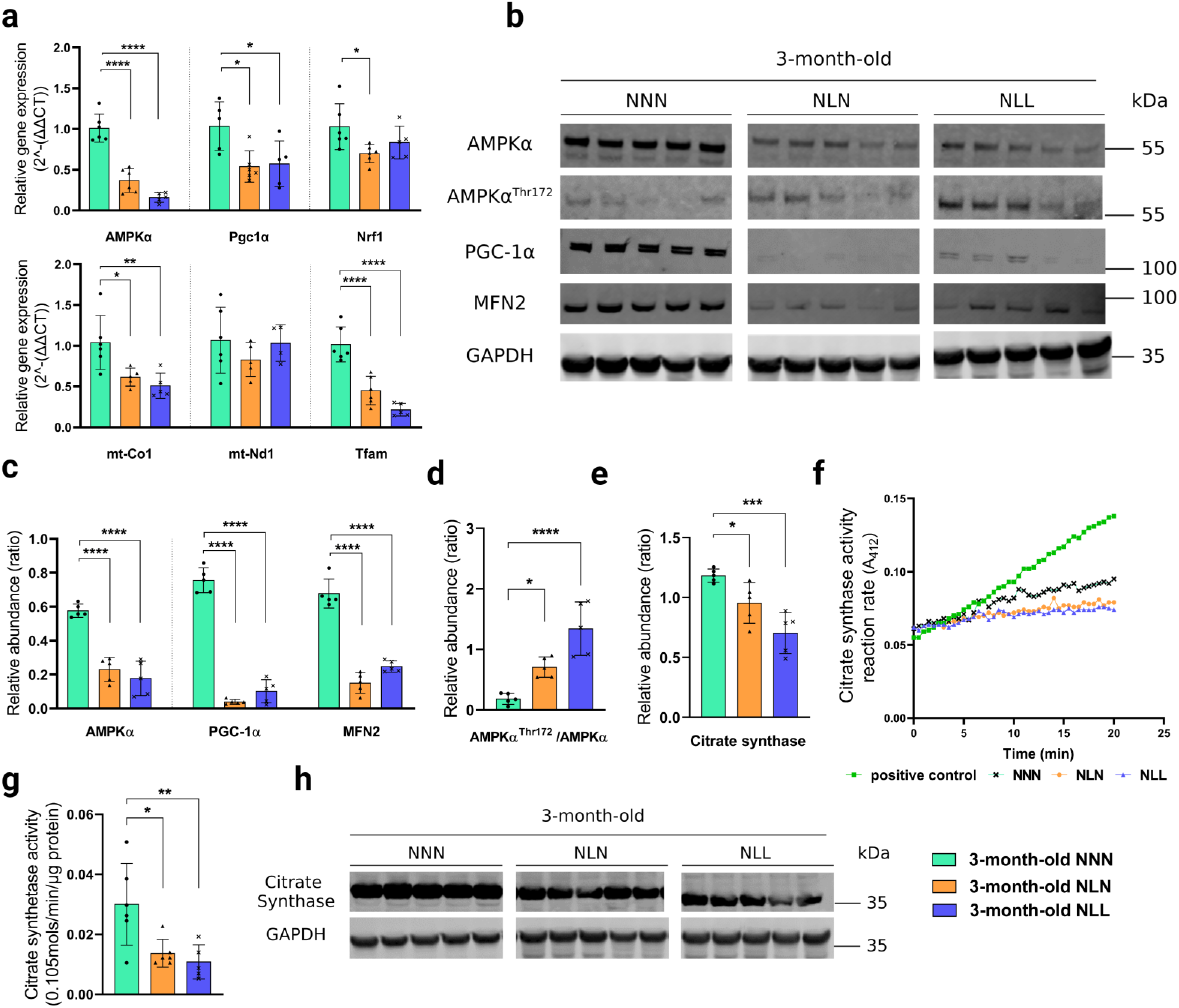
Lifelong protein restriction impairs AMPK signalling and mitochondrial homeostasis in skeletal muscle. (**a**) Relative gene expression levels of AMPKα, Peroxisome proliferator-activated receptor gamma coactivator 1-alpha (Pgc-1α), Nucleus respiratory factor 1 (Nrf-1), mitochondrially encoded cytochrome c oxidase subunit I (mt-Co1), Mitochondrially Encoded NADH: Ubiquinone Oxidoreductase Core Subunit 1 (mt-Nd1), and Transcription Factor A, Mitochondrial (Tfam) in GAS muscle from 3-month-old mice *n=5-*6. **(b, c, d)** Western blot analysis of (b) AMPKα, AMPKα^Thr172^, PGC-1α, and Mitofusin 2 (MFN2) in GAS muscle from NNN, NLN, and NLL mice Samples were run on the same gel and images were cropped only for the purpose of this figure. GAPDH or ponceau staining were used as loading control. Source data provided as a Source data file. *n=5.* (c) Quantification of AMPKα, PGC-1α, and MFN2 protein abundance levels. (d) Ratio of AMPKα^Thr172^/AMPKα protein abundance. *n=5.* **(f)** Representative graph of citrate synthase reaction rate (A_412_) in positive control and GAS muscle. **(g)** Citrate synthase activity (0.105*μM/min/μg protein*) in GAS muscle from each experimental group. The assays were performed in duplicate for each sample. *n=5-6.* **(h, e)** Western blot analysis of (h) citrate synthase in GAS muscle from 3-month-old NNN, NLN, and NLL mice. *n=5.* (e) Quantification of citrate synthase protein abundance levels in GAS muscle Samples were run on the same gel and images were cropped only for the purpose of this figure. GAPDH was used as loading control. Source data provided as a Source data file. *n=5.* p<0.05, ⃰ ⃰ p<0.01, ⃰ ⃰ ⃰ p<0.001, ⃰⃰⃰ ⃰ ⃰ ⃰ p<0.0001, ordinary one-way ANOVA with a Dunnett’s multiple comparisons test.

**Fig. 7.**
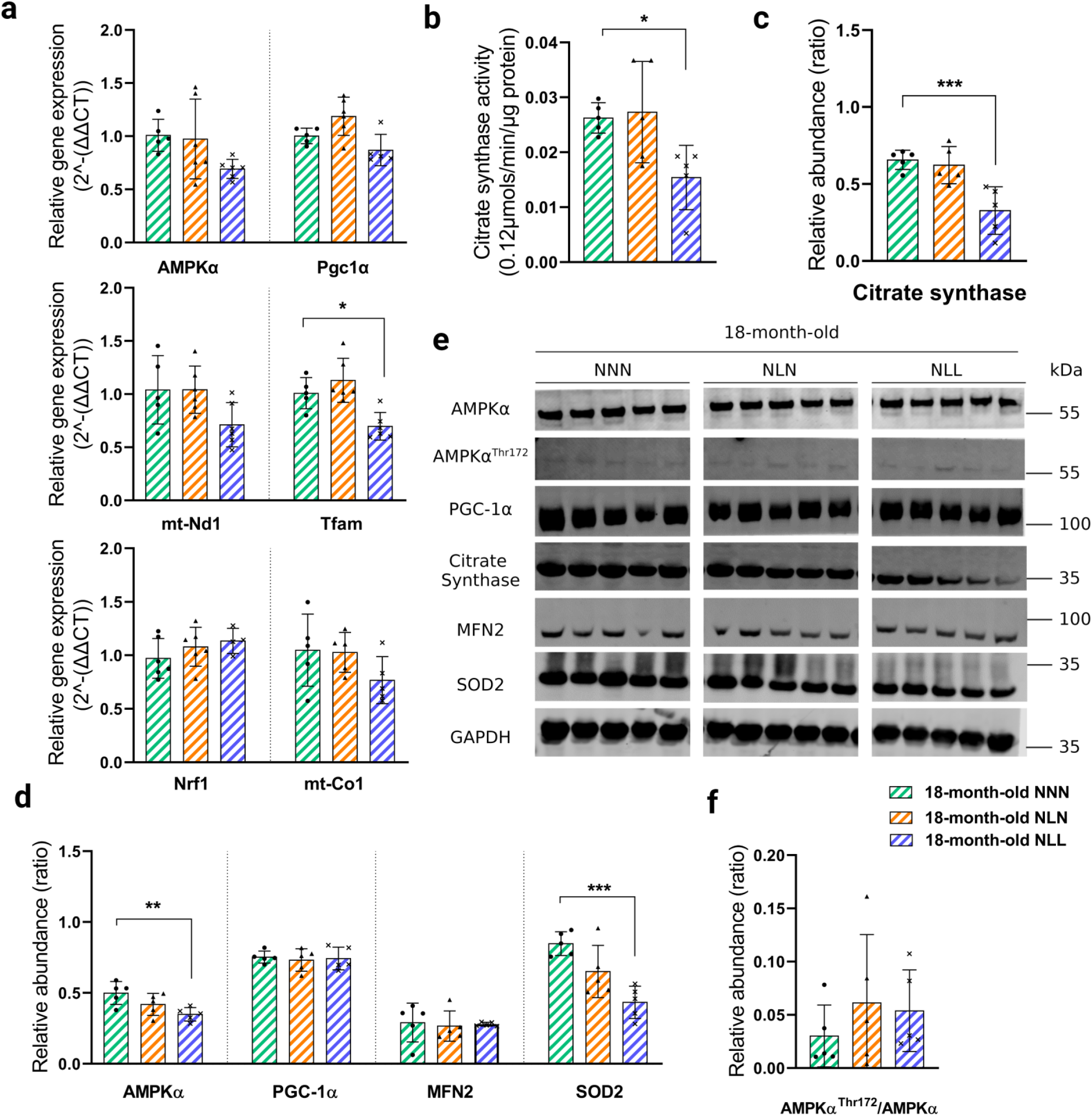
The detrimental effects of lactational protein restriction on the AMPK signalling pathway and mitochondrial biogenesis can be corrected by weaning onto a normal protein content diet in later life. (**a**) Relative gene expression levels of AMPKα, PGC-1α, NRF-1, mt-Co1, mt-Nd1, and TFAM in GAS muscle from 18-month-old mice *n=5-*6. **(b)** Citrate synthase activity (*μM/min/120μg protein*) in GAS muscle from each experimental group. The assays were performed in duplicate for each sample. *n=5-6* (c) Quantification of citrate synthase protein abundance levels in GAS muscle. *n=5*. **(d)** Quantification AMPKα, PGC-1α, MFN2, and SOD2 protein abundance levels. *n=5.* (e) Samples were run on the same gel and images were cropped only for the purpose of this figure. GAPDH or ponceau staining were used as loading control. Source data provided as a Source data file. *n=5.* (f) Ratio of AMPKαThr172/AMPKα protein abundance. *n=5.* ⃰ p<0.05, ⃰ ⃰ p<0.01, ⃰ ⃰ ⃰ p<0.001 ordinary one-way ANOVA with a Dunnett’s multiple comparisons test.

**Fig. 8.**
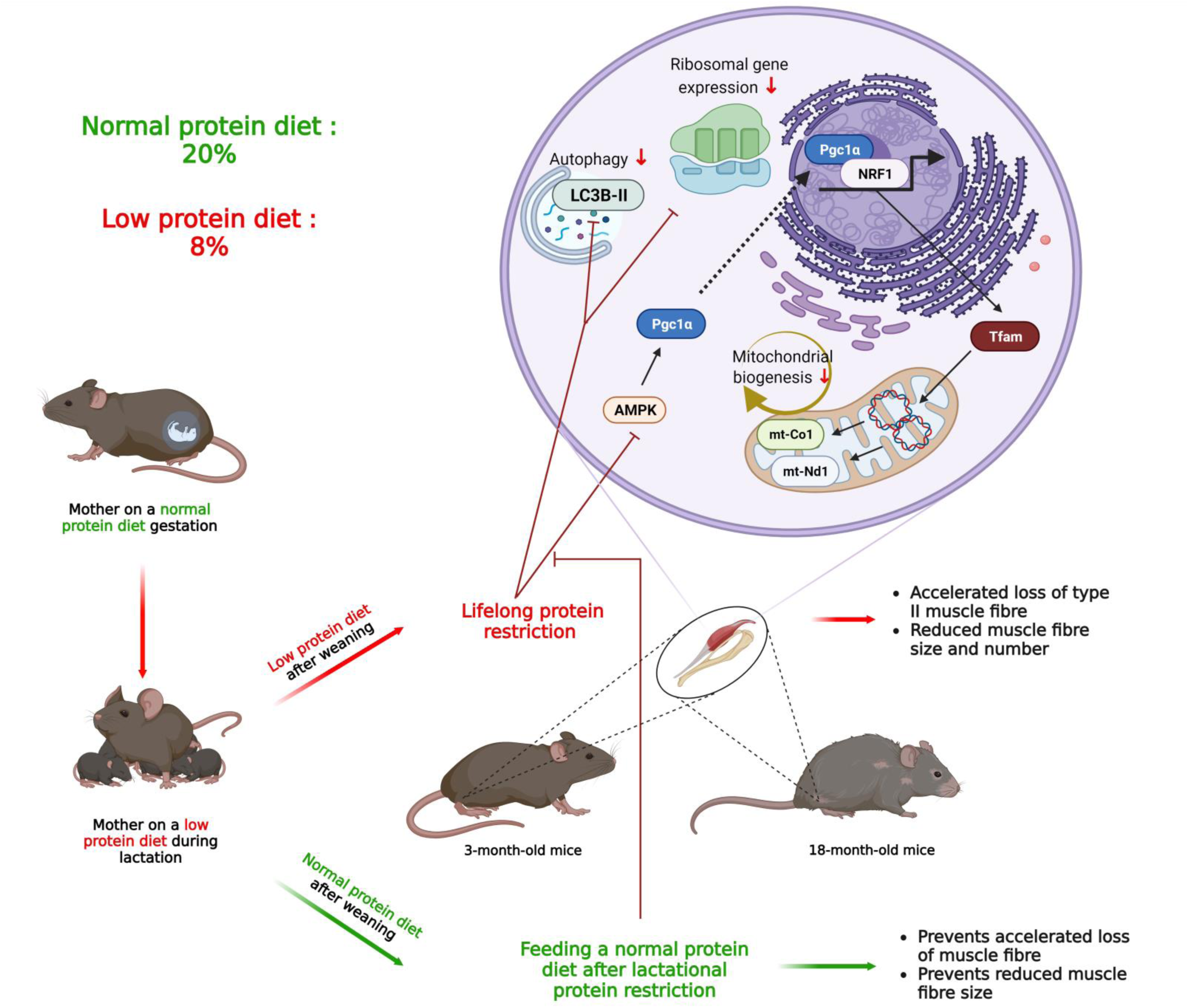
Effect of lactational protein restriction followed by either a low protein diet or normal protein diet after weaning, on skeletal muscle in later life.

We further supported this mechanism by investigating Peroxisome proliferator-activated receptor gamma coactivator 1-alpha (PGC1α), which is a transcription factor that induces mitochondrial biogenesis by activating transcriptional factors such as Nucleus Respirator Factor 1 (NRF1) [32]. We found a dramatic reduction in PGC1α protein abundance (Fig. 6b, 6c) and a downregulation at the gene expression level of Pgc1α in NLL mice at 3 months of age (Fig. 6a). There was no significant difference in Nrf1 gene expression level between 3-month-old NLL and NNN mice (Fig. 6a). Gene expression and protein content of PGC1α and Nrf1 gene expression levels did not differ between the groups at 18 months of age.

As mitochondrial biogenesis depends on the nucleus for transcriptional factors to increase the expression of mitochondrial protein-encoding genes, we next assessed gene expression levels of Transcription Factor A, Mitochondrial (Tfam), which regulates the expression of mitochondrially-encoded NADH: Ubiquinone Oxidoreductase Core Subunit 1 (mt-Nd1) and mitochondrially encoded cytochrome c oxidase subunit 1 (mt-Co1) [29]. RNA-seq showed slight reduction in Tfam, mt-Co1, and Mt-Nd1 expression in GAS muscle from 21-day-old NL mice (Supplementary Data 1). However, Tfam expression was significantly reduced in GAS from NLL mice (Fig. 6a, Fig. 7a). At 3 months of age, the expression of the mt-Co1 gene was significantly lower in NLL mice compared to NNN mice (Fig. 6a). However, there was no significant difference in mt-Co1 expression level between NLL and NNN mice at 18 months of age (Fig. 7a), nor was there a significant difference in mt-Nd1 expression level at either time points (Fig. 6a, Fig. 7a). These findings were in line with lifelong protein restriction resulting in a reduction in mitochondrial biogenesis.

Impaired AMPK signalling and reduced mitochondrial biogenesis led us to hypothesize that other contributors to mitochondrial homeostasis, such as mitochondrial fusion and mitophagy would also be affected in GAS muscle from NLL mice. Western blot analyses revealed that there was a significant reduction in the protein content of Mitofusin-2 (MFN2) (Fig. 6b, 6c) and UNC-51-like kinase 1 (ULK1) (Fig. 5d, 5e) in GAS muscle from NLL mice at 3 months of age. In contrast, our findings in 18-month-old mice showed no significant difference in MFN2 (Fig. 7d, 7e) and ULK1 (Fig. 5f, 5g) protein abundance in NLL mice compared to the control group.

Mitochondrial reactive oxygen species (ROS) production in muscle cells are important in redox signalling and damage mitochondria in a variety of pathologies [33]. Hence, we next investigated the antioxidant enzymes, superoxide dismutase (SOD2) and peroxiredoxin 3 (PRDX3) to determine the effects of impaired AMPK signalling and mitochondrial homeostasis on mitochondrial antioxidant defence. At 21 days of age, RNA-seq showed no difference in Sod2 and Prx3 gene expression in NL mice in comparison to the control group (Supplementary Data 1). There was a significant downregulation of SOD2 at 3 and 18 months after feeding a low-protein diet following lactational protein restriction (Supplementary Fig. 1a, 1c). Additionally, gene expression of Sod2 was significantly reduced in 3-month-old NLL mice compared to the control group (Supplementary Fig. 1b). Furthermore, we found a significant decrease in gene expression and protein content of PRDX3 in GAS muscle from NLL mice at 3 months of age (Supplementary Fig. 1a, 1b, 1c). Thus, feeding a lifelong low-protein diet following lactational protein restriction may dysregulate mitochondrial antioxidant defence.

### The detrimental effects of lactational protein restriction on ribosomal gene expression, autophagy, AMPK pathway, and mitochondrial homeostasis were corrected by weaning onto a normal protein content diet

Our RNA-seq data revealed that there was an upregulation in the expression level of genes involved in histone phosphorylation in 21-day-old NL mice (Supplementary Table 3). The long-lasting detrimental effects of gestational maternal manipulations are often explained through changes in the epigenetic landscape, which is driven by modifications of cytosine bases and histones [34–36]. Hence, this led us to hypothesise that changes seen in skeletal muscle from 21-day-old mice may be long-lasting through epigenetic modifications into later life. To explore this, we investigated ribosomal gene expression, autophagy, the AMPK signalling, and mitochondrial homeostasis in muscles of mice that were fed on a normal protein diet after lactational protein restriction (NLN), both at 3 months and 18 months of age.

There was a significant increase in gene expression levels of Rpl36a and Rpl38 in skeletal muscle from NLN mice at 3 months of age (Fig. 5b). This is consistent with NLN mice exhibiting an accelerated catch-up growth following lactational protein restriction-induced reduction in body weight. However, we did not observe any significant differences in the expression levels of Rpl36a, Rpl38, and Rps7 between 18-month-old NLN and NNN mice (Fig. 5c). We also analysed autophagy markers in mice that were fed on a normal-protein diet after lactational protein restriction. At 3 months of age, p62 protein abundance was significantly lower in NLN mice compared to the control group (Fig. 5d, 5e), but there was no significant difference between the two groups at 18 months of age (Fig. 5f, 5g). Moreover, LC3B-II/LC3B-I ratio did not show any significant difference between the two groups at either time point (Fig. 5d, 5e, 5f, and 5g).

A significant reduction in AMPKα, Pgc1α, Nrf1, mt-Co1, and Tfam gene expression levels (Fig. 6a) and protein content of AMPKα, PGC1α, and MFN2 (Fig. 6b, 6c) were observed in GAS muscle from 3-month-old NLN compared to the control mice. This was accompanied by a significant reduction in both activity and abundance of citrate synthase in 3-month-old NLN mice (Fig. 6e, 6f, 6g, 6h). Additionally, protein abundance of ULK1 (Fig. 5d, 5e), SOD2, PRDX3 (Supplementary Fig. 1a, 1c) and gene expression level of Sod2 and Prdx3 (Supplementary Fig. 1b) were substantially reduced in NLN mice at 3 months of age.

Next, we investigated whether lactational protein restriction would have similar effects on AMPK signalling pathway and mitochondrial homeostasis at 18 months of age as they had at 3 months of age. Strikingly, there was no significant difference between 18-month-old NLN and NNN mice in terms of the expression level of genes and content of proteins involved in AMPK signalling and mitochondrial homeostasis (Fig. 7). Notably, the activity and abundance of citrate synthase were similar in these mice at 18 months of age (Fig. 7b, 7c, 7e).

## Discussion

Sarcopenia is a progressive and widespread skeletal muscle disorder associated with adverse outcomes including frailty, mortality, and falls [37]. Therefore, it is important to acquire a comprehensive understanding of how sarcopenia develops. It has been reported that the principal cause of sarcopenia is loss of muscle fibre number and fibre atrophy – notably among type II fibres [38–40]. Protein deficiency is recognized as a widespread and persistent global issue due to its multifactorial causes, including limited access to expensive food sources containing essential amino acids, hindering the optimal growth and development of the offspring [7]. In this study, we assessed the effects of lifelong protein restriction on GAS and SOL skeletal muscles and our results demonstrated that feeding a lifelong low-protein diet following lactational protein restriction accelerated the loss of type IIa fibres and reduced muscle fibre size in later life. Consistently, our group previously reported that a postnatal low-protein diet (5%, compared to normal protein diet 17-20%) reduced maximum force generation in extensor digitorum longus (EDL) muscle at 3 months of age in mice [41]. Another animal study has reported that rats born to mothers fed a low protein diet (8%) and maintained on an 8% protein diet in postnatal life had a decreased cross-sectional area and distorted myofibers in their GAS muscle at 16 weeks of age [42]. Furthermore, our previous research demonstrated that long-term protein restriction led to a consistent decrease in tibialis anterior (TA) muscle weight throughout the lifespan of male mice [43]. Together, this data suggested that lifelong protein restriction may subsequently contribute to the development of sarcopenia, but further studies are required to follow this up after 18 months of age when the sarcopenic phenotype is fully developed.

Our data indicated that lactational protein restriction lead to a significant upregulation of ribosomal gene expression in the GAS skeletal muscle in the offspring. Since protein deficiency during lactation resulted in limited growth in the offspring, these findings suggest that the observed increase in ribosomal mRNA expression serves as a compensatory response to the growth restriction. Similar to these findings, our group previously demonstrated that lactational protein restriction also upregulated the expression of small nucleolar RNAs (snoRNAs), such as U3, in skeletal muscle. snoRNAs facilitate multiple roles in ribosomes biogenesis including 2′-O-methylation of ribosomal (r)RNA and pseudouridylation [44]. Hence, given the vital role of U3 in the maturation of ribosomal rRNAs, it has been proposed that the upregulation of U3 expression may also act as an adaptive mechanism in response to protein restriction during lactation [26]. Moreover, feeding a lifelong low-protein diet after lactational protein restriction reduced ribosomal gene expression and autophagy in GAS muscle in later life. This decrease in autophagy was associated with the accelerated loss of skeletal muscle fibres and reduced muscle fibre size later in life. Although autophagy flux is increased during the process of muscle wasting [45], there is accumulating evidence to suggest that impaired autophagy may contribute to age-related muscle loss [46–48]. For example, a rat study has reported an age-related decline in autophagic degradation in skeletal muscle [48]. Together, it can be inferred that lifelong protein restriction leads to an acceleration in skeletal muscle loss by impairing autophagy and ribosomal gene expression, thus compromising proteostasis.

We also demonstrated that feeding a low-protein diet following lactational protein restriction disturbed AMPK signalling and impaired mitochondrial homeostasis and mitochondrial antioxidant defence in later life. This suggested that mitochondria play a key role in the accelerated skeletal muscle loss phenotype observed in mice fed a low-protein diet following lactational protein restriction. Consistent with our findings, a loss in muscle mass has been related to a decline in mitochondrial biogenesis [49, 50], while PGC1α-TFAM activation alleviates skeletal muscle atrophy [51]. In addition, Mfn2 deletion resulted in muscle atrophy and deletion of both Mfn1 and Mfn2 decreased muscle performance in mice [52, 53], whereas stimulating mitophagy can counteract age-related skeletal muscle loss [54–56]. It has been reported that mitochondria-targeted antioxidants may prevent immobilisation-induced muscle atrophy [57, 58]. Our study offers mechanistic insight into lifelong protein restriction-induced skeletal muscle loss, which could be a targeting strategy for studies investigating muscle wasting.

It is well-established that maternal undernutrition has a great effect on the metabolic health outcome of the offspring [8]. In the present study, we found that lactational protein restriction impaired ribosomal protein gene expression, regulation of mitochondrial ATP synthesis coupled electron transport, the transition between fast and slow fibre and skeletal muscle fibre development. While previous studies have reported long-lasting adverse effects of maternal protein restriction on offspring in later life via epigenetic modifications, these studies mostly focused on the influence of gestational protein restriction on the offspring [26, 41, 59]. Longitudinal studies following women who were pregnant during the Dutch Famine of World War II have reported the impact of maternal nutrition on offspring development and their susceptibility to diseases in humans [60]. In contrast, we provide evidence that the detrimental effects of lactational protein restriction in skeletal muscle can be corrected by weaning onto a normal protein diet in later life in mice. Similarly, our group previously reported that male and female mice fed a normal protein diet following lactational protein restriction exhibited a compensatory growth response and had similar survival rate to the control groups [43]. In addition, Ozanne and Hales, 2004 described that iso-caloric protein restriction during gestation shortened the lifespan in mice, whereas the offspring nursed by mothers on a low-protein diet during lactation lived longer than the control mice [15]. Collectively, these findings suggest that lactational protein restriction may not have long-lasting detrimental effects on skeletal muscle, but could even have beneficial effects when followed by a normal protein diet starting at weaning. Translation of our findings into human health has the potential to revolutionize the guidelines for appropriate maternal nutritional diet during pregnancy and lactation. Further studies are needed to examine the importance of the timing of maternal nutritional programming in mediating the long-term adverse effects on the offspring.

We showed that lifelong low protein diet after lactational protein restriction had adverse effects on skeletal muscle at 18 months of age. However, reduced dietary consumption of protein is linked with increased metabolic health and longer life span in flies, rats, mice, and humans [61–65]. Richardson and colleagues (2021) reported that lifelong consumption of low amino-acids (AA) diet significantly increased the lifespan of C57BL/6J male mice [66]. Moreover, lifelong dietary branched-chain amino acids (BCAA) restriction decreased the mammalian target of rapamycin (mTOR) signalling in skeletal muscle and the frailty in mice [66]. Key differences from our study include protein restriction during lactation, dietary design, and the restriction of only BCAAs. In their study, Richardson and colleagues examined the effects of a 7% protein diet starting at weaning or the specific restriction of BCAAs by 67% [66]. Our study involved the restriction of dietary protein intake in lactating mothers and the subsequent feeding of their offspring with a low-protein diet beginning at weaning. This highlights the importance of maternal dietary protein intake during lactation. To comprehensively evaluate the differences, further research is required, focusing on a direct comparison between a low protein diet administered during lactation and a low protein diet initiated after weaning.

In summary, lactational protein restriction hinders skeletal muscle growth by impairing mitochondrial and ribosomal homeostasis. While there is a substantial amount of work indicating the adverse effects of maternal malnutrition during gestation on the offspring later in life, for the first time we have shown that at least some of the detrimental effects of lactational protein restriction seen in skeletal muscle can be attenuated by weaning onto a normal content protein diet. However, lifelong protein restriction accelerated skeletal muscle loss and reduced muscle fibre size in later life, and this was associated with an impairment in the AMPK signalling pathway and the dysregulation of mitochondrial homoeostasis and proteostasis. Hence, one should consider the AMPK-PGC1α-mitochondria axis and proteostasis as potent factors in protein restriction-driven skeletal muscle loss.

## Methods Animals

B6.Cg-Tg(Thy1-YFP)16Jrs/J mice (Jackson Laboratory; stock number 003709), were used for this study. Individual vented cages were utilized to house the mice with a fixed light cycle (21 ± 2°C, 12-hour light/dark cycle). Ethical approval was obtained from the University of Liverpool Animal Welfare Ethical Review Committee (AWERB), and UK Animals (Scientific Procedures) Act 1986 regulations were followed for all experimental protocols for the handling and use of laboratory animals.

## Experimental design

Two weeks prior to mating, nulliparous female mice were fed either a low-protein diet (L, 8% crude protein; Special Diet Services, UK; code 824248) or a normal-protein diet (N, 20% crude protein; Special Diet Services, UK; code 824226), both fed *ad libitum*. Age-matched male mice on a normal-protein diet were used for mating. When the female mice were pregnant, they were kept on the same diet. The male offspring from mothers on normal-protein diet during gestation were used for this study. Within 24 hours of birth, the new-born male litters were cross-fostered to mothers fed either a normal-protein diet or a low-protein diet, in order to create Normal-Normal (NN) or Normal-Low (N-L) groups of mice, respectively. Suckling pup numbers were kept the same for all animals during lactation (n=6 pups). Following weaning, NN mice were fed a normal protein diet generating Normal-Normal-Normal (NNN or control, n=6) mice (Fig. 1a). In order to understand the effects of lactational protein restriction and to establish if this could be corrected by feeding a normal diet from weaning, NL mice were weaned onto either a normal protein diet, creating Normal-Low-Normal (NLN, n=6), (Fig. 1a). Moreover, NL mice were fed a low-protein diet to investigate the effects of lifelong protein restriction, generation Normal-Low-Low (NLL, n=6), (Fig. 1a). Mice were fed *ad libitum* food and water. At 3-month-old or 18-month-old of age, mice were sacrificed. Body weights were recorded immediately, prior to dissection. GAS and SOL muscles were carefully dissected and weighed. Muscle tissues were stored at –80°C until analysis and prepared for the experiments as described below.

## RNA isolation, library preparation and sequencing

Frozen GAS skeletal muscles of 21-day-old mice (n = 6) were ground using a mortar and pestle in liquid nitrogen. Total RNA from frozen GAS muscle was extracted and purified by RNeasy mini kits with on-column DNase treatment (Qiagen, Manchester, UK) following manufacturer’s instructions. Preparation of dual-indexed, strand-specific RNA-seq library from submitted total RNA was performed using the NEBNext polyA selection and Ultra II Directional RNA library preparation kits (New England Biolabs, UK) as described previously [67]. Total RNA integrity was confirmed by an Agilent 2100 Bioanalyzer (Agilent Technologies, Santa Clara, CA, USA) with RNA Integrity Number (RIN)>7. RNA-seq carried out by Illumina NovaSeq using S1 chemistry (paired-end, 2×150bp sequencing, generating an estimated 650 million clusters per lane). Number of reads obtained by RNA-seq were provided in Supplementary Figure 4. Sequencing was performed at the Centre for Genomic Research, University of Liverpool (https://www.liverpool.ac.uk/genomic-research/). The raw RNA-seq data have been deposited in NCBI Gene Expression Omnibus (GEO) database under accession number GSE235750.

## Data processing and bioinformatic analyses

The raw Fastq files are trimmed by the Centre for Genomic Research, University of Liverpool for the presence of Illumina adapter sequences using Cutadapt version 1.2.1 [68]. Any reads which match the adapter sequence for 3 bp or more are trimmed. The reads are further trimmed using Sickle version 1.200 with a minimum window quality score of 20. Reads shorter than 15 bp after trimming were removed. The processed reads were then pseudo-aligned to the Ensemble release *mus musculus* transcriptomes v96 with Kallisto quant version 0.46.1 to get the transcripts length and abundance estimates [69] The output from these tools was converted count by tximport (TXI) version 1.26.1. in R version 4.2.2 [70]. An average of 41.96 million total counts for NN and 45.32 million total counts for NL were generated following alignment (Fig. 2a). Differentially expression and statistical analyses were carried out by DESeq2 version 1.38.3. and PCA plots were visualised by plotPCA function in ggplot2 version 3.4.1 in R. We used the following cutoffs:|Log2 Fold Change| (|Log2FC|)>0.3 and adjusted p-values<0.1 to compare gene expressions. To generate the heatmap, we selected the 130 genes which had |Log2FC|>1 and adjusted p-values<0.1 using the ‘pheatmap’ library version 1.0.12 in R.

To further analyse potential biological pathways and perform pathway enrichment analysis, we utilized ShinnyGO version 0.77 [71], which incorporates Gene Ontology (GO), Kyoto Encyclopaedia of Genes and Genomes (KEGG), and Reactome databases. All differentially expressed genes were inputted into ShinnyGO, with protein coding genes from mouse species selected as the background. FDR<0.05 was used for the fold enrichment and GO terms. To calculate fold enrichment, the percentage of differentially expressed genes within a pathway is divided by the corresponding percentage of such genes in the background. STRING version 11.5 were used to create network images [72].

## Gene expression analyses

Total RNA was extracted from ground GAS or SOL muscles using TRIzol (Thermo Fisher Scientific, Waltham, MA, USA) and cleaned up using RNeasy mini kits (Qiagen, 74104) following manufacturer’s instructions. A DNAse digestion step was performed on extracted RNA using RNase-Free DNase Set (Qiagen, Manchester, UK) to eliminate the risk of genomic contamination. cDNA was then synthesized from 1000 ng mRNA using the High-Capacity cDNA Reverse Transcription Kit (Applied Biosystems, Warrington, UK). Quantitative PCR was performed using Meridian Bioscience SensiMix SYBR Hi-ROX Kit (Scientific Laboratory Supplies, Nottingham, UK) in 20 µL reaction volumes. Data were analysed using Rotor-Gene Q Series software (Qiagen, Manchester, UK). Expression levels were normalized to Glyceraldehyde 3-phosphate dehydrogenase (Gapdh) expression at 3 months of age and 18S rRNA expression at 21 days and 18 months of age, as the expression levels of these reference genes did not differ between groups at the given age (Supplementary Figure 3). The delta-delta Ct method (2^−ΔΔCt^) were performed to calculate the relative fold gene expression [73]. All primers were validated prior to gene expression analyses. Primers are listed Supplementary Table 1.

## Antibodies

Primary antibodies used for Western blotting are listed as follows, the dilutions are indicated in the list. Anti-PGC1 alpha (ab191838, dilution 1:1000), Anti-NOX2/gp91phox (ab129068, 1:500), Anti-GCLC (ab190685, 1/500), Anti-PRDX3 (ab73349, 1:1000), Anti-p62 (ab91526, 1:1000), Anti-Citrate synthase (ab129095, 1:1000), Anti-GAPDH (ab8245, 1:2000) from Abcam (Cambridge, UK); Anti-AMPKα (#2532, 1:400), Anti-Phospho-AMPKα^Thr172^ (#2535, 1:400), Anti-MFN2 (#9482, 1:500), Anti-ULK1 (#8054, 1:200), Anti-Phospho-ULK1^Ser555^ (#5869, 1:250), Anti-LC3B (#43566, 1:250), from Cell Signalling (Cell Signaling Technology, Danvers, MA, USA); Anti-SOD2 (ADI-SOD-111, 1:2000) from Enzo Life Sciences (Farmingdale, New York, USA). Primary antibodies Anti-MYH I (BA-D5, 1:200), Anti-MYH IIb (BF-F3, 1:250), and Anti-MYH IIa (SC-71, 1:250) from Developmental Studies Hybridoma Bank (Iowa, USA) and Wheat Germ Agglutinin (WGA), CF®405S Conjugate (29027-1, 1:200, Generon, Slough, UK) for membrane staining were used for immunohistochemistry.

## Western blotting

Total protein was extracted from a part of the grounded GAS muscles. RIPA Lysis Buffer (Sigma, Poole, UK) containing Pierce Protease and Phosphatase Inhibitor Mini Tablets, Ethylenediaminetetraacetic acid (EDTA)-free (Thermo Fisher Scientific, Waltham, MA, USA). The samples were homogenized and sonicated for 30 seconds twice, and then centrifuged for 10 minutes at 14,000 x g at 4°C. Supernatant was collected and total protein content determined with a BCA Protein Assay (Sigma, Poole, UK). 30 or 45 μg protein extract were electrophoresed and separated on NuPAGE™ 4 to 12%, Bis-Tris, 1.0–1.5 mm, Mini Protein Gels (Invitrogen, Renfrewshire, UK) and transferred to a polyvinylidene fluoride (PVDF) membrane. GAPDH (ab8245, 1:2000) or ponceau staining (Sigma, Poole, UK) were used as loading control. Membranes were blocked with 5% bovine serum albumin (BSA) (Sigma, Poole, UK) in tris-buffered saline (TBS) for an hour and were cut to probe with primary antibodies which were optimised prior to experiments. Pre-cut blots were used for the antibody treatment overnight. After incubating with a fluorescence secondary antibody, blots were imaged on a the Licory Odissey CLx (Licor, Bad Homburg, Germany) and band densities were analysed using ImageJ. Results were normalised to the loading control.

## Histological analyses

SOL muscles were carefully dissected and orientated vertically on cork discs. Optimal Cutting Temperature (OCT – Thermo Fisher Scientific, Waltham, MA, USA) embedded muscles were immersed in liquid nitrogen cooled isopentane. 8 μm thick sections were cut using a Leica 1850 (Leica Biosystems, Newcastle upon Tyne, UK) for histological analyses. For Myosin Heavy Chain (MYH) staining, sections were air dried for 30 minutes and fixed with 100% ice-cold acetone. Cryosections were blocked using Mouse On Mouse (M.O.M) (Vector Laboratories Ltd., Peterborough, UK) blocking with 5% Goat Serum in Phosphate Buffered Saline (PBS) and incubated with primary antibodies overnight, and subsequently with secondary fluorescent antibodies (Invitrogen) for an hour at room temperature. Sections were mounted with VECTASHIELD Antifade Mounting Medium (Vector Laboratories Ltd., Peterborough, UK), and images captured using a Zeiss LSM 800 confocal microscope (Zeiss, Oberkochen, Germany). Images were analysed using ImageJ (U.S. National Institutes of Health, USA) and MyoVision software (University of Kentucky, USA).

## Citrate synthase assay

GAS muscles were powdered in liquid nitrogen and the resulting powder was lysed in extraction buffer (250 mM Sucrose (Sigma-Aldrich, Gillingham, Dorset, UK), 10 mM Trizma Base (Sigma-Aldrich, Gillingham, Dorset, UK), and 1 mM EGTA (Sigma-Aldrich, Gillingham, Dorset, UK)). Samples were then sonicated for 30 seconds twice on ice and centrifuged for 10 minutes at 12,000 x g at 4°C. Bradford assay (Thermo Fisher Scientific, Waltham, MA, USA) was used to quantify the protein content of supernatants. MitoCheck® Citrate Synthase Activity Assay kit (Cayman Scientific, Ann Arbor, MI, USA) was utilized to measure citrate synthase reaction rate by recording 412 nm absorbance every 30 seconds for a total of 20 minutes. Recordings were plotted against time. Subsequently, the slope of the linear curve was calculated and used to quantify the reaction.

## Statistical analyses

In all cases, *n* refers to the number of litters, and only one male per litter was used for any one outcome. GraphPad Prism (Dotmatics, Boston, Massachusetts, USA) software version 8.0.2 were used to analyse the data, which are expressed as mean ± standard deviation (mean ± SD) with n=4-6. Statistical comparisons were performed using ordinary one-way ANOVA with a Dunnett’s multiple comparisons test, considering NN/NNN as the control group. A p<0.05 level of confidence was considered for statistical significance. Data was checked for normal distribution with Shapiro-Wilk normality test.

## Acknowledgements

We thank the Biomedical Services Unit (BSU), University of Liverpool for excellent animal care throughout the project. The authors wish to thank Karen Guerrero Vazquez for providing technical help with data analysis.

## Funding

I.K., A.V., and K.G-W disclose support for the research of this work from Biotechnology and Biological Sciences Research Council (grant number BBSRC; BB/P008429/1). U.E. discloses support for the research of this work from the Turkish Embassy in London. M.J.J. and M.J.P. disclose support the research of this work from the MRC-Versus Arthritis Centre for Integrated research into Musculoskeletal Ageing (CIMA). The authors disclose support for publication of this work from the University of Liverpool.

## Contributions

I.K., K.G-W, and A.V. conceived and designed the experiments with input from S.E.O; U.E carried out the experiments; U.E., M.A., I.K., M.J.P., M.J.J., K.G-W, and A.V. analysed and elaborated data; U.E. and G.P.V. performed the bioinformatic analysis; I.K., M.J.P., S.E.O, M.J.J., K.G-W, and A.V. provided critical comments to the manuscript; U.E wrote the manuscript.

## Ethic declarations Conflict of interest

The authors declare that they have no competing interests.

## Notes

https://www.ncbi.nlm.nih.gov/geo/query/acc.cgi?acc=GSE235750

### Competing Interest Statement

The authors have declared no competing interest.

