## Supplementary figures and images for "Lifelong dietary protein restriction accelerates skeletal muscle loss and reduces muscle fibre size by impairing proteostasis and mitochondrial homeostasis"

### Source Data

#### Slide 1
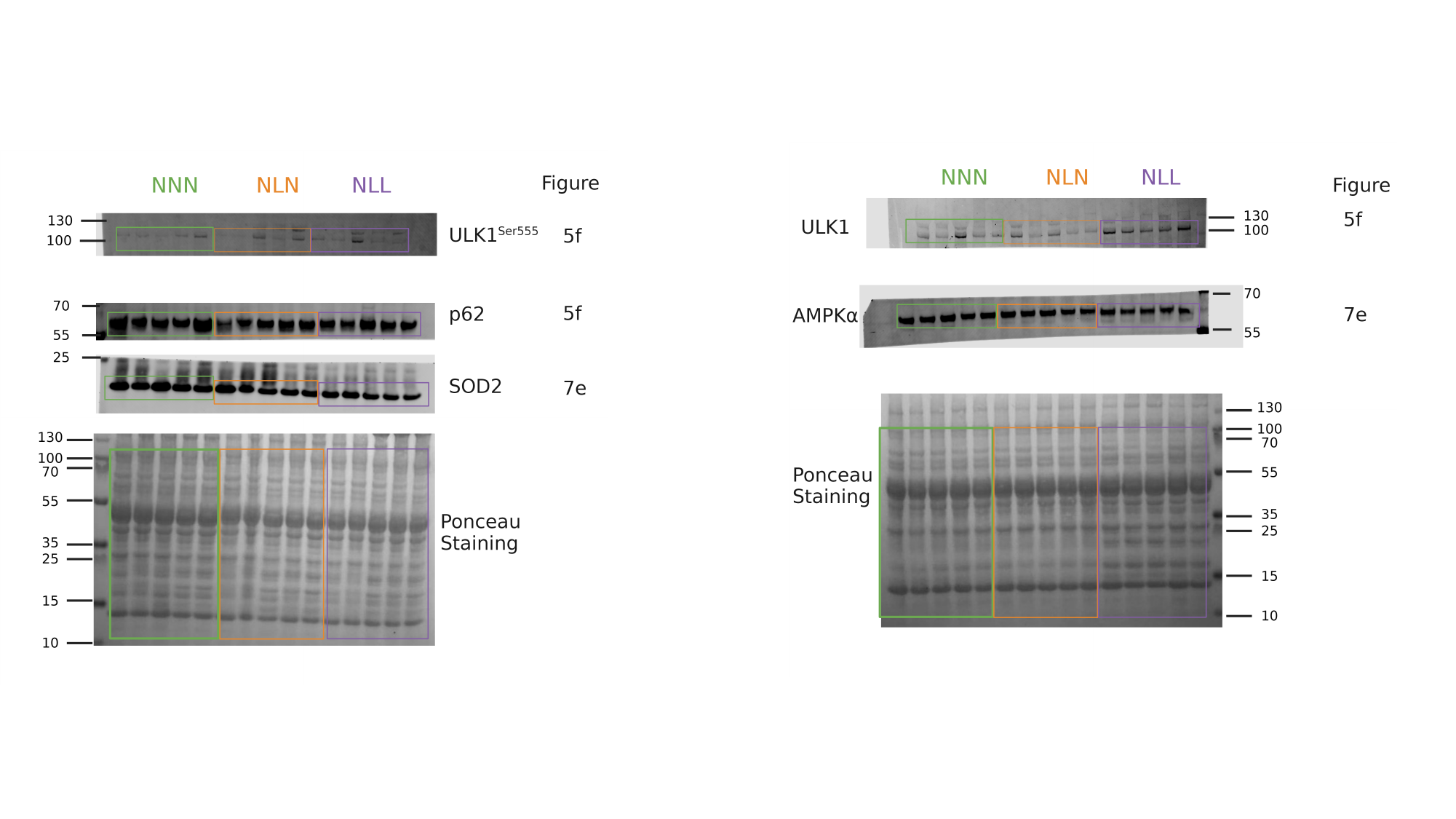

#### Slide 2
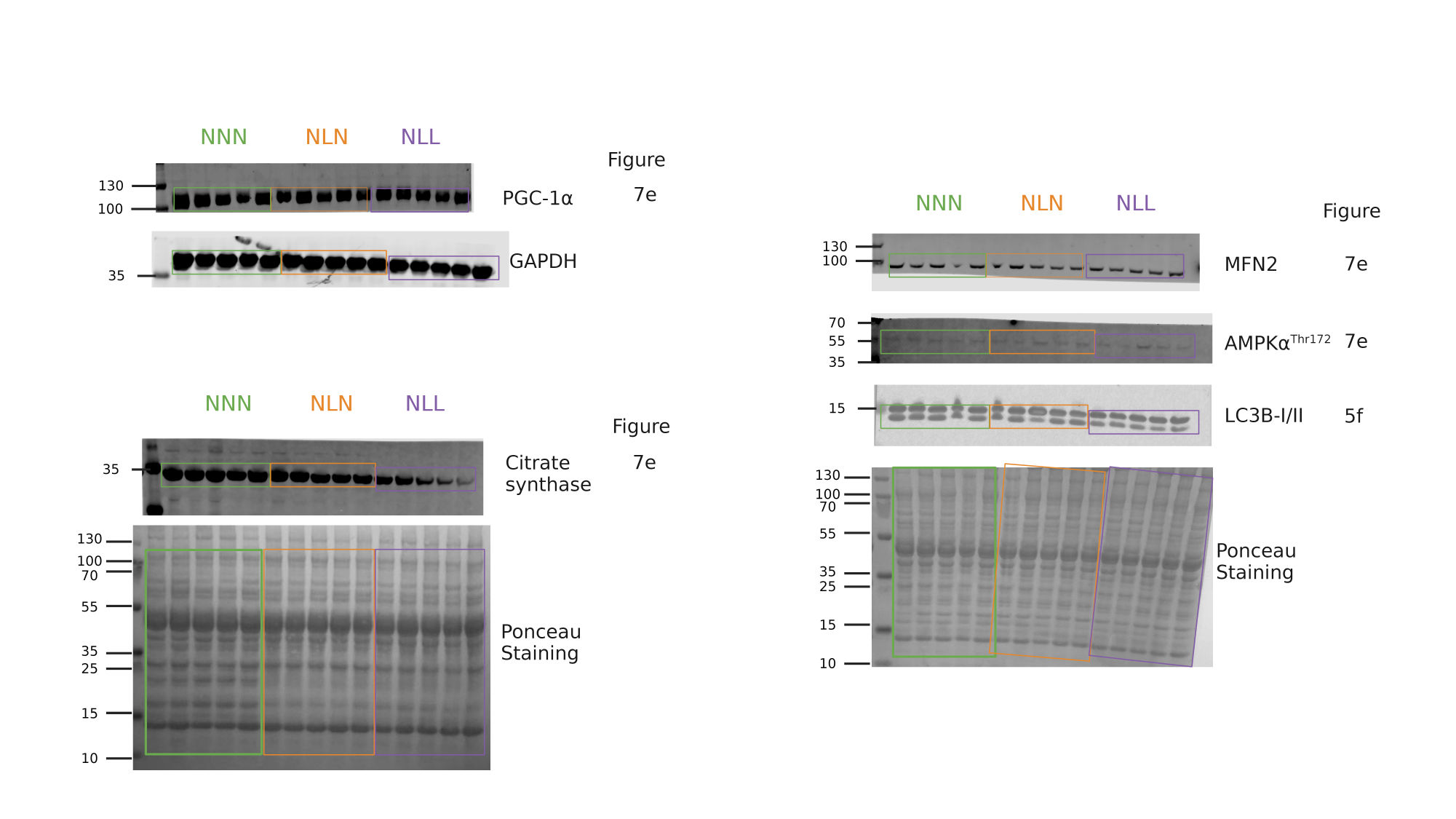

#### Slide 3
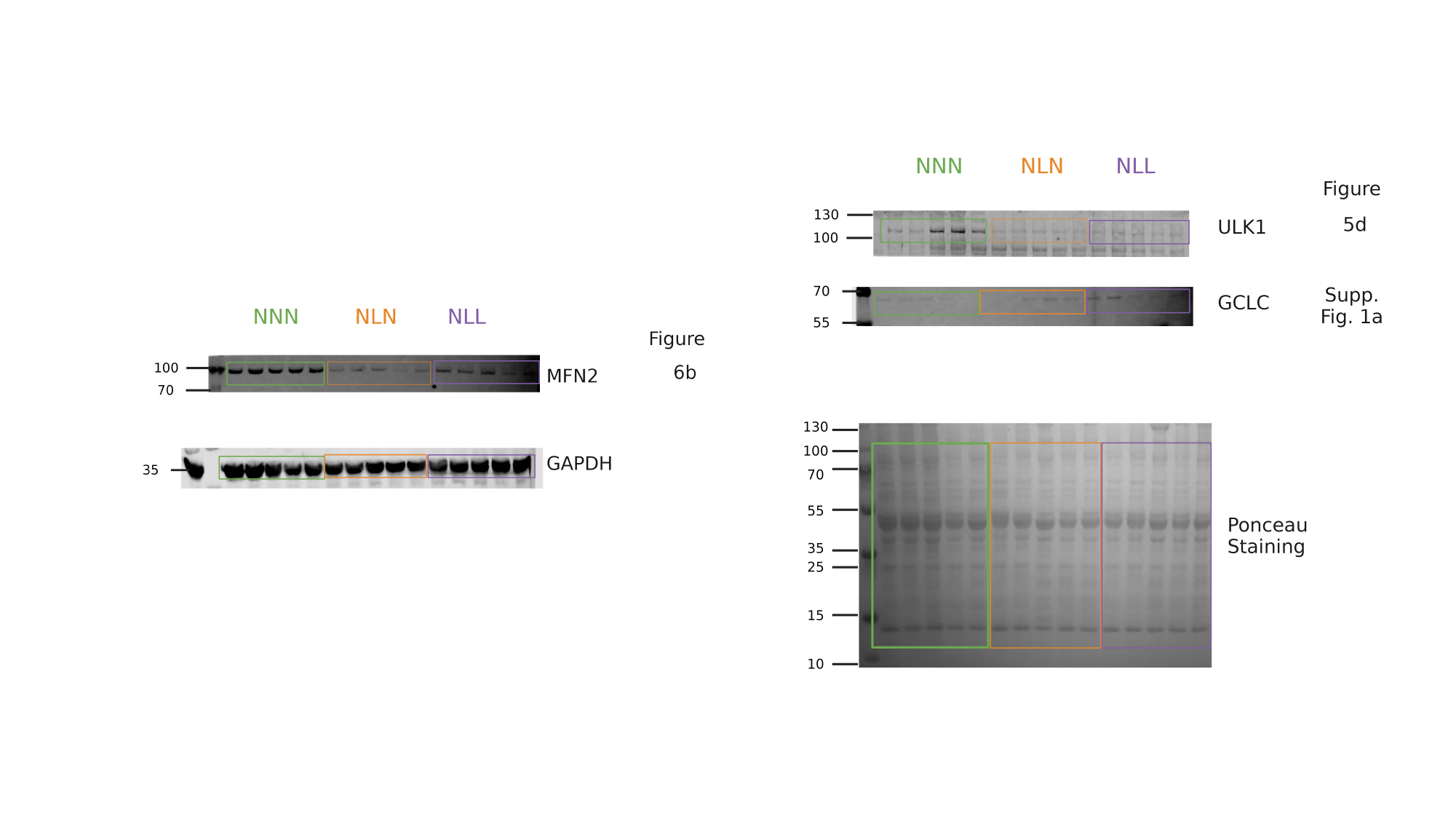

#### Slide 4
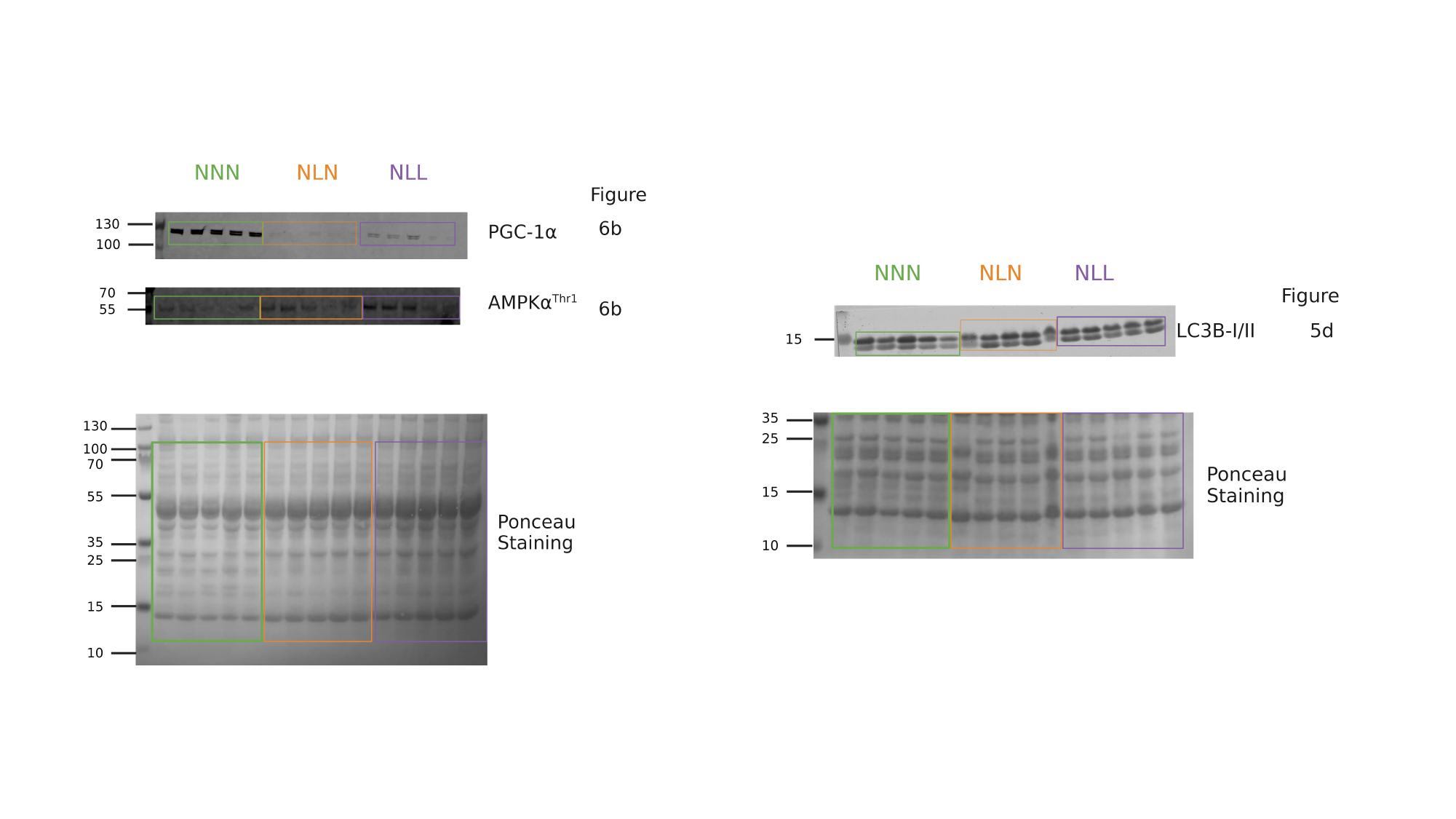

#### Slide 5
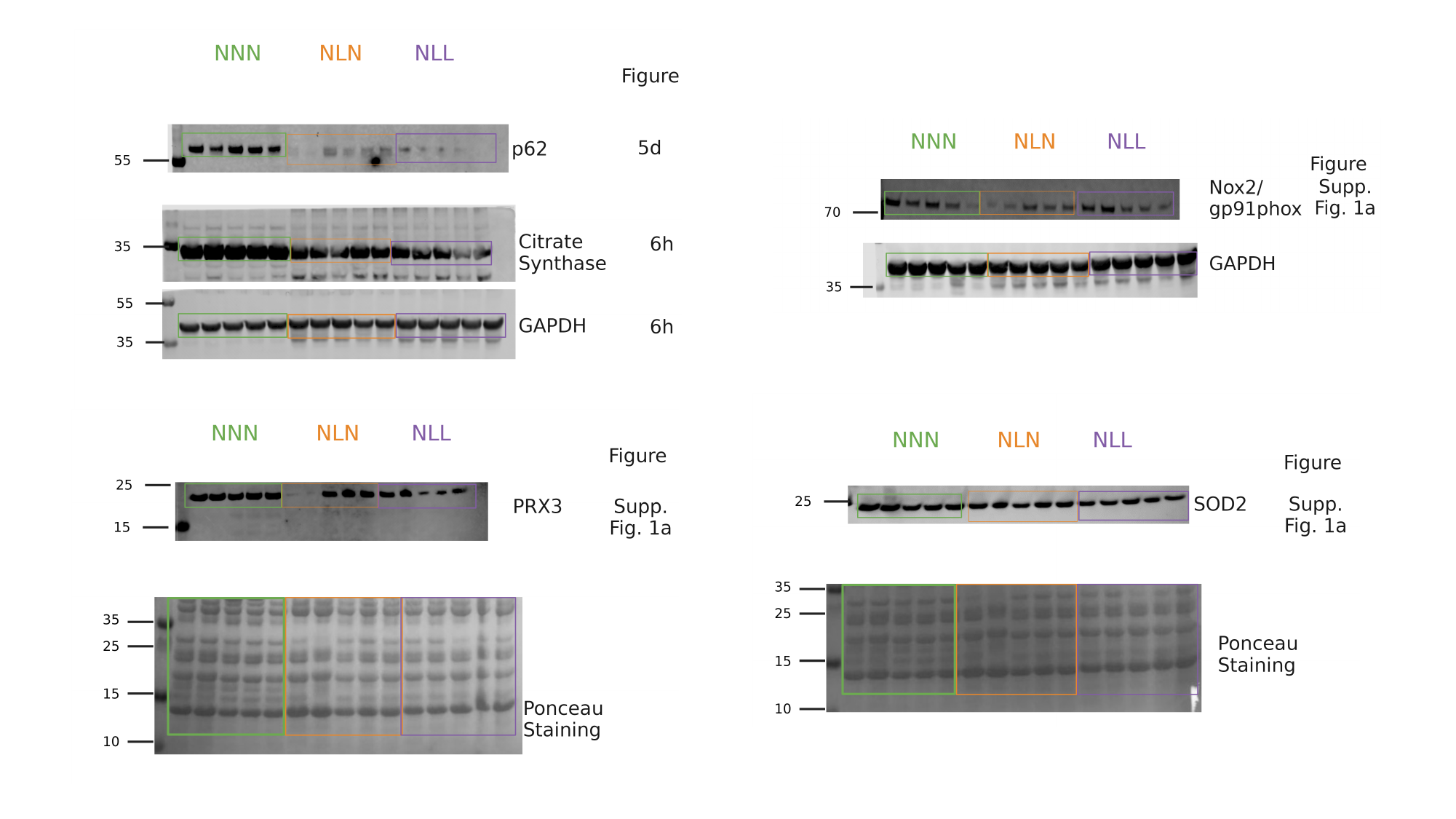

#### Slide 6
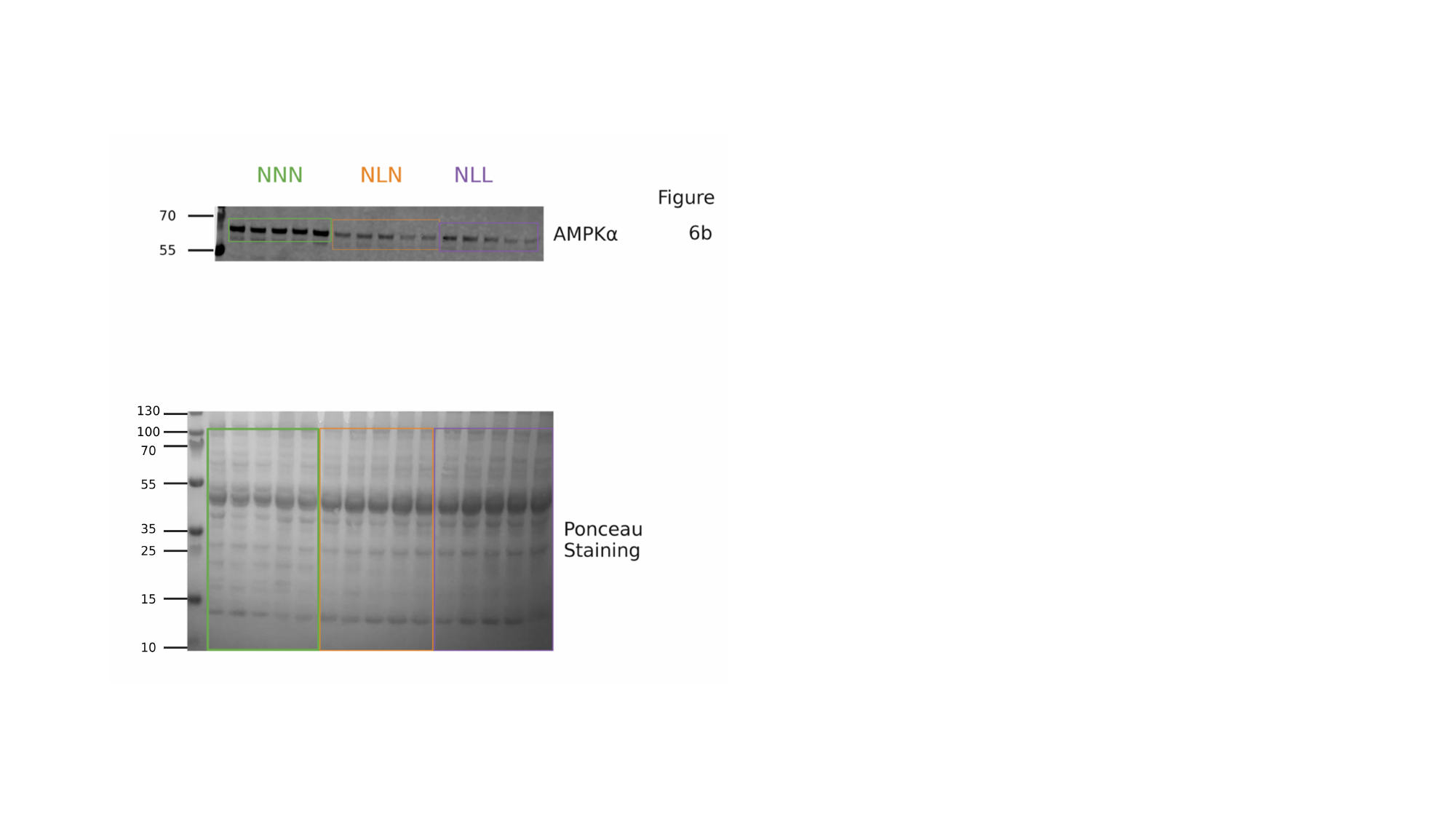
