## Supplementary Figures for "Lifelong dietary protein restriction accelerates skeletal muscle loss and reduces muscle fibre size by impairing proteostasis and mitochondrial homeostasis"

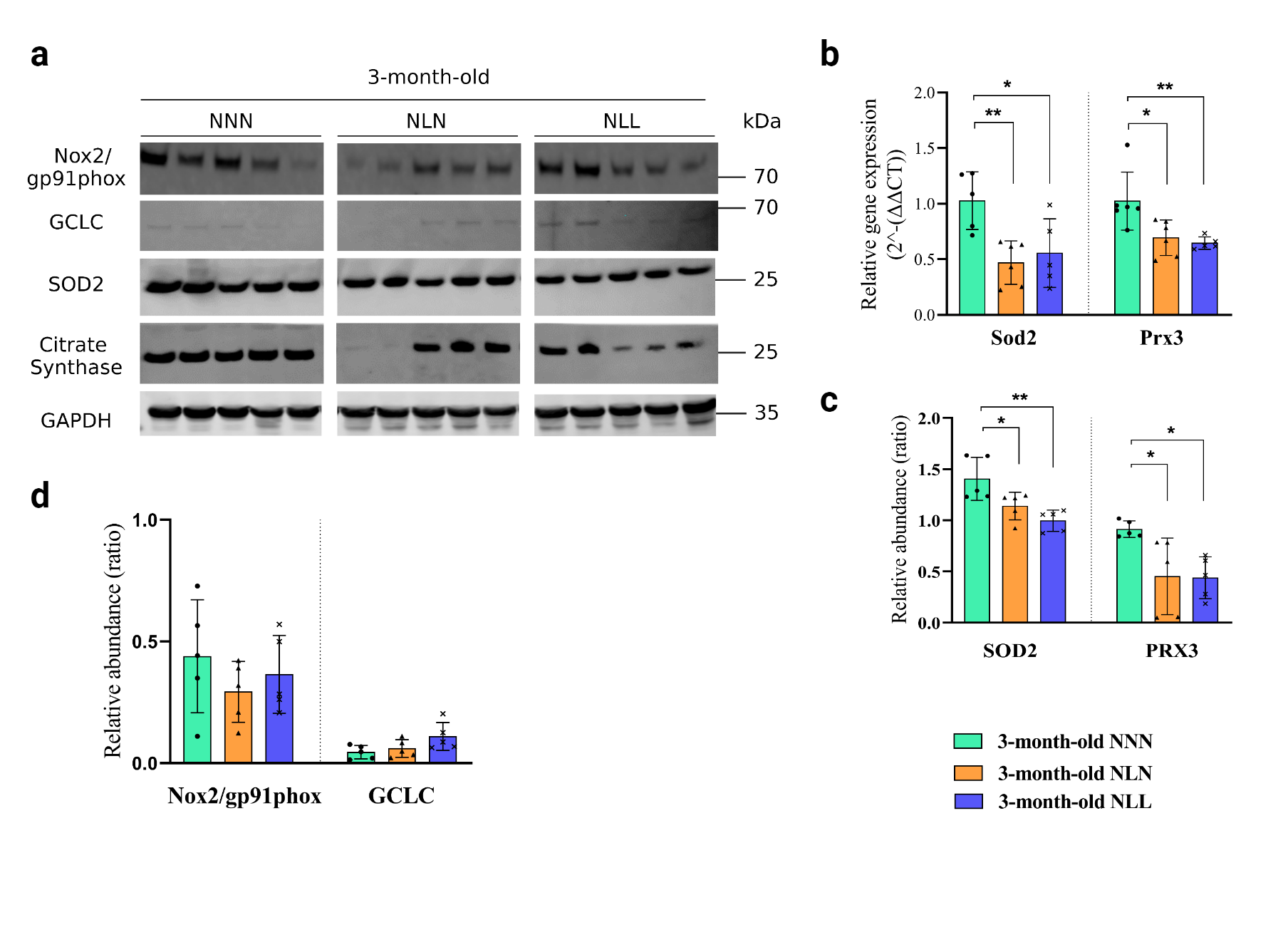

**Supplementary Figure 1. The effects of prolonged protein restriction or feeding a normal protein diet after lactational protein restriction on oxidative stress markers and antioxidants in skeletal mice from adult mice. (a, c, d)** Western blot analysis of (a) Nox2/gp91phox, GCLC, SOD2, and PRX3 in GAS muscle from NNN, NLN, and NLL mice. Samples were run on the same gel and images were cropped only for the purpose of this figure. GAPDH or ponceau staining were used as loading control. Source data provided as a Source data file. *n=5.* (c) Quantification of SOD2 and PRX3 protein abundance levels. (d) Quantification of Nox2/gp91phox, p40phox, and GCLC protein abundance levels. **(b)** Relative gene expression levels of Sod2 and Prx3 in GAS muscle from 3-month-old mice *n=5-*6. Results are expressed as the mean ± standard deviation (mean ± SD). ⃰ p<0.05, ⃰ ⃰ p<0.01, ⃰ ⃰ ⃰ p<0.001, ⃰⃰⃰ ⃰ ⃰ ⃰ p<0.0001. Statistical comparisons were performed using ordinary one-way ANOVA with a Dunnett’s multiple comparisons test, considering NNN as the control group.

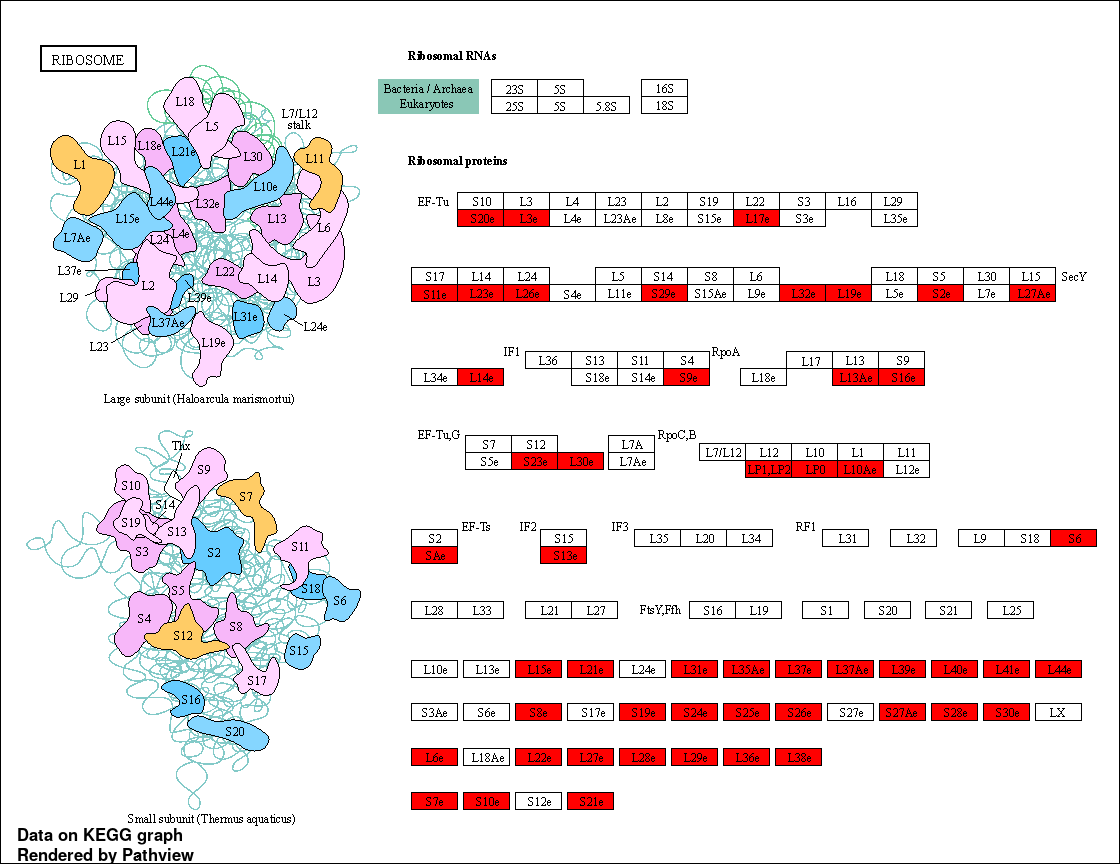

**Supplementary Figure 2. Illustration of the differential expressed ribosomal genes (shown in red) in GAS muscle from 21-day-old lactational protein-restricted mice.**

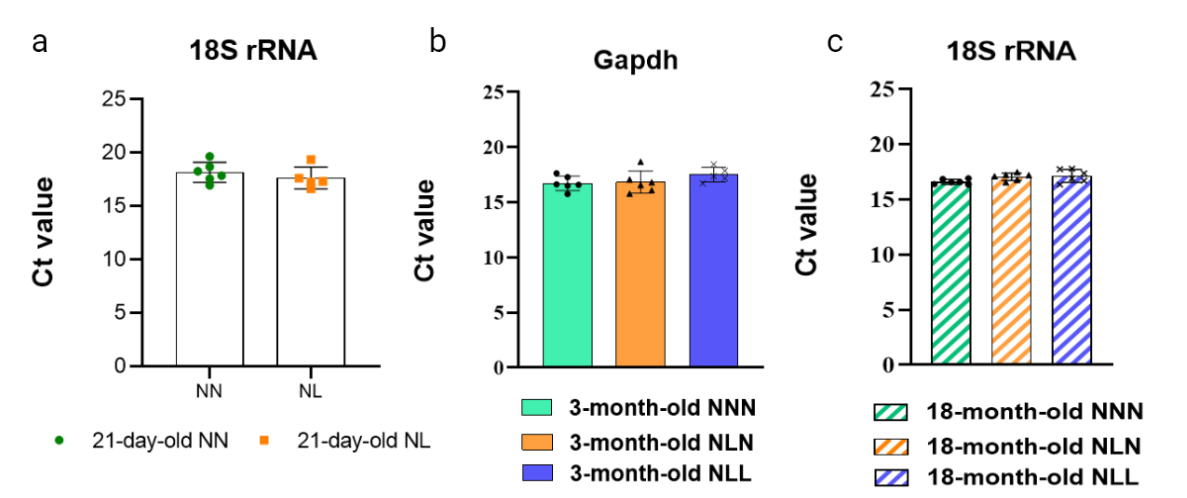

**Supplementary Figure 3. Expression level of Glyceraldehyde 3-phosphate dehydrogenase (Gapdh) and 18S rRNA reference genes di not differ between groups. (a)** 18s Ct value in GAS skeletal muscle from 21-day-old NN and NL mice. **(b)** Gapdh Ct value in GAS skeletal muscle from 3-month-old NNN, NLN, and NLL mice. **(c)** 18s Ct value in GAS skeletal muscle from 18-month-old NNN, NLN, and NLL mice. Results are expressed as the mean ± standard deviation (mean ± SD). ⃰ p<0.05, ⃰ ⃰ p<0.01, ⃰ ⃰ ⃰ p<0.001, ⃰⃰⃰ ⃰ ⃰ ⃰ p<0.0001. Statistical comparisons were performed using ordinary one-way ANOVA with a Dunnett’s multiple comparisons test, considering NNN as the control group.

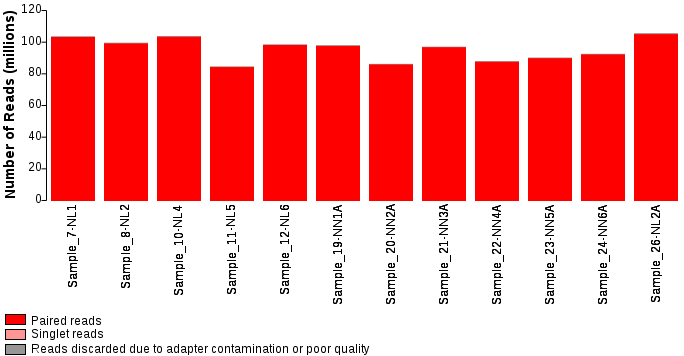

**Supplementary Figure 4. Number of reads obtained by RNA-seq per sample.**

**Supplementary Table 1** List of primers. AMP-activated protein kinase alpha 1 (AMPKa1), Glyceraldehyde 3-phosphate dehydrogenase (Gapdh), mitochondrially encoded cytochrome c oxidase I (mt-Co1), mitochondrially encoded NADH: Ubiquinone Oxidoreductase Core Subunit 1 (mt-Nd1), Myosin Heavy Chain (Myh), Nucleus Respirator Factor 1 (Nrf-1), Peroxisome proliferator-activated receptor gamma coactivator 1-alpha (Pgc1α), peroxiredoxin 3 (Prdx3), ribosomal protein L36A (Rpl36a), ribosomal protein L38 (Rpl38), ribosomal protein S7 (Rps7), Transcription Factor A, Mitochondrial (Tfam), superoxide dismutase (Sod2), and 18S rRNA.

| *Gene* | Forward sequence | Reverse sequence | Exon-exon spanning | Primer efficiency | Validation |
| --- | --- | --- | --- | --- | --- |
| *AMPKa1* | ggaaagtgaaggtgggcaa | atagctctcctccagagacat | - | 102.4% | + |
| *Gapdh* | catcactgccacccagaagactg | atgccagtgagcttcccgttcag | - | 91.3% | + |
| *mt-Co1* | cccagatatagcattcccacg | actgttcatcctgttcctgc | - | 101.7% | + |
| *mt-Nd1* | ctagcagaaacaaaccgggc | ccggctgcgtattctacgtt | - | 109.03% | + |
| *MyhI* | ggagcgcaagtttgtcataagt | ctcaagctgctcagcaatctattt | + | 104.0% | + |
| *MyhIIa* | ctccaaggaccctcttatttccc | actgctgaactcacagaccc | + | 94.1% | + |
| *MyhIIb* | tgtgtgtccttcagcattccc | gaggcaatcaggaaccttcgg | + | 101.1% | + |
| *Nrf1* | gtcgctcatccaggttggta | gatggtcatttcaccgccct | + | 93.1% | + |
| *Pgc1a* | gacaggtgccttcagttcac | caaccagagcagcacactcta | + | 95.2% | + |
| *Prdx3* | gtatctccgcctatcgtgcc | catgacgagcaaccgagga | + | 86.7% | + |
| *Rpl36a* | caagaagagaaagggccaagt | tattccctcccaaacgaacaa | + | 89.6% | + |
| *Rpl38* | ttccccgttctcttcggttc | aattttccgaggcatggcga | + | 97.9% | + |
| *Rps7* | atgaactccgatctcaaggc | caccaacttcgatttccttgg | + | 100.7% | + |
| *Tfam* | cataggcaccgtattgcgtg | tcggaatacagacaagactgataga | + | 89.8% | + |
| *Sod2* | ttaacgcgcagatcatgca | ggtggcgttgagattgttca | - | 106.3% | + |
| *18S* | ggaaagcagacatcgacctca | agttctccagccctcttggt | - | 95.2% | + |
